# FOXO-regulated Deaf1 controls muscle regeneration through autophagy

**DOI:** 10.1101/2024.01.12.575306

**Authors:** Kah Yong Goh, Wen Xing Lee, Sze Mun Choy, Gopal Krishnan Priyadarshini, Kenon Chua, Qian Hui Tan, Shin Yi Low, Hui San Chin, Chee Seng Wong, Shu-Yi Huang, Nai Yang Fu, Jun Nishiyama, Nathan Harmston, Hong-Wen Tang

## Abstract

The commonality between various muscle diseases is the loss of muscle mass, function, and regeneration, which severely restricts mobility and impairs the quality of life. With muscle stem cells (MuSCs) playing a key role in facilitating muscle repair, targeting regulators of muscle regeneration has been shown to be a promising therapeutic approach to repair muscles. However, the underlying molecular mechanisms driving muscle regeneration are complex and poorly understood. Here, we identified a new regulator of muscle regeneration, Deformed epidermal autoregulatory factor 1 (Deaf1) - a transcriptional factor downstream of FOXO signaling. We showed that Deaf1 is transcriptionally repressed by FOXOs and that Deaf1 targets to PI3KC3 and Atg16l1 promoter regions and suppresses their expressions. *Deaf1* depletion therefore induces autophagy, which in turn blocks MuSC survival and differentiation. In contrast, *Deaf1* overexpression inactivates autophagy in MuSCs, leading to increased protein aggregation and cell death. Interestingly, *Deaf1* depletion and overexpression both lead to defects in muscle regeneration, highlighting the importance of fine tuning Deaf1-regulated autophagy during muscle regeneration. We further showed that *Deaf1* expression is altered in aging and cachectic MuSCs. Remarkably, manipulation of *Deaf1* expression can attenuate muscle atrophy and restore muscle regeneration in aged mice or mice with cachectic cancers. Together, our findings unveil an evolutionarily conserved role for Deaf1 in muscle regeneration, providing insights into the development of new therapeutic strategies against muscle atrophy.

## Introduction

Skeletal muscle accounts for approximately 40–50% of total body weight and plays key roles in movement, thermogenesis, and maintenance of energy homeostasis [1]. With the constant need for adaptability and the ability to acclimatize to physical demands such as growth and hypertrophy training, it is vital that skeletal muscles are able to regenerate from injury or regular wear and tear [2, 3]. This regenerative ability is primarily dependent on a population of MuSCs, also known as satellite cells, which are located underneath the basal lamina of the myofiber [3]. Under resting conditions, MuSCs are generally characterized by the expression of the myogenic transcription factor, paired box 7 (Pax7) and are maintained in a quiescent state [4]. During muscle growth or in response to injury, cytokines and growth factors activate MuSCs, driving them out of quiescence and into proliferation, which leads to the generation of myoblasts. Myoblasts are myogenic progenitors which differentiate and fuse together to form multinucleated myofibers, repairing muscle injuries and promoting growth [4, 5]. This muscle regeneration process is regulated by transcription cascades, including multiple myogenic regulatory factors, such as MyoD, Myf5, Myogenin and MRF4 [4].

Declination of muscle regenerative functions has been linked to various muscle diseases such as sarcopenia, which is a common age-related skeletal muscle disorder featuring the loss of muscle mass accompanied by reduced levels of MuSCs [6]. Sarcopenic patients are known to exhibit decreased MuSCs and this decline has also observed in muscles of old mice [6, 7]. Furthermore, MuSCs isolated from aged mice and injected into young muscle displayed defects in self-renewal, expansion, and myogenic differentiation, suggesting that cell-autonomous changes in aged MuSCs disrupts muscle regeneration even in young milieu [8, 9]. Several alterations in multiple signal transduction cascades in aged MuSCs have been identified, including the inactivation of forkhead box O (FOXO) transcription factors [10], and aberrant activation of Jak-Stat3 [11] and the stress-associated p38a/b mitogen-activated protein kinase (MAPK) pathways [12]. In addition, proteotoxicity caused by reduced levels of autophagy is another key contributing factor [3, 13, 14]. Autophagy is a cell survival mechanism that enables cells to adapt to stress by means of degradation and recycling of long-lived proteins or damaged organelles and is regulated by multiple autophagic regulators [15-19]. Autophagic inactivation in aging MuSCs causes the accumulation of damaged mitochondria, which induces ROS and DNA damage and ultimately leads to senescence and apoptosis [13], suggesting that autophagy-regulated proteostasis is essential for preservation of stemness.

Cancer cachexia is a tumor-induced syndrome characterized by a loss of skeletal muscle weight and impaired MuSC functions [20]. It was found that the proliferation and differentiation of MuSCs are compromised during cancer cachexia [21, 22]. Apoptosis in myoblasts has also been observed in cancer cachexia [23]. In contrast to sarcopenia, the expressions of FOXOs were increased in skeletal muscles under cachectic conditions [24, 25]. Inhibition of FOXOs’ activities in cachectic mice increased the levels of MyoD, a myogenic factor, and decreased Myostatin, leading to a marked increase in MuSC proliferation as well as skeletal muscle mass [24, 25]. These studies suggest that FOXOs in MuSCs play critical roles in cancer-induced muscle wasting. Contrary to sarcopenia, autophagic activation in MuSCs contributes to muscle wasting in cancer cachexia [26]. Despite similar defects observed in both sarcopenia and cancer cachexia such as muscle loss and reduced muscle regeneration, the underlying molecular mechanism leading to each could potentially be different.

Stem cell therapy utilized for muscle diseases has been showing substantial advancements with its ability to repair and promote muscle regeneration in several animal experiments [6, 27]. To enhance the efficacy and safety of stem cell therapy, elucidation of mechanisms underpinning its role is required. To identify novel MuSC regulators, we utilized a cell lineage tracing approach and performed a RNAi screen in *Drosophila* MuSCs. We identified Deformed epidermal autoregulatory factor 1 (Deaf1) as a novel regulator of muscle regeneration. Surprisingly, we found that both overexpression and knockdown of *Deaf1* inhibited *Drosophila* muscle regeneration through the regulation of autophagy. Whilst *Deaf1-RNAi* upregulates autophagy and activates cell death, overexpression of *Deaf1* inhibits autophagy, leading to accumulation of protein aggregates and apoptosis in MuSCs. These phenomena were also observed in C2C12 myoblasts and in primary MuSCs isolated from mouse muscle. We further identified FOXO transcription factors as upstream regulators of Deaf1. FOXO1 and FOXO3 bind to the Deaf1 promoter, inhibiting Deaf1 expression, and in turn activates transcription of autophagy-related genes, *Atg16l1* and *PI3KC3*, as demonstrated by ChIP-seq analyses. Importantly, both aging and cachectic MuSCs display altered *Deaf1* expression. Pharmaceutical or adeno-associated virus serotype 9 (AAV9)-mediated modulation of *Deaf1* expression relieved muscle regeneration defects induced by aging or cancer, highlighting the key roles of Deaf1 in sarcopenia and cancer cachexia. Together, we conclude that Deaf1 is a key regulator of autophagy during muscle regeneration and is a potential druggable target for sarcopenia and cancer cachexia.

## Results

### A genetic screen identifies *Drosophila* Deaf1 as a regulator of MuSCs

To gain insights into the modulators of skeletal MuSCs, we performed a RNAi screen in *Drosophila* utilizing cell lineage analysis. The GAL4 Technique for Real-time and Clonal Expression (G-TRACE) system contains five elements: *Zfh1-Gal4, UAS-RFP* fluorescent protein, *UAS-Flipase*, *Ubi-FRT-STOP-FRT-EGFP*, and *tubGal80^ts^* (Figure 1A). It has been reported that *Zfh1* is specifically expressed in MuSCs [28-30]. In this system, the enhancer elements of *Zfh1* controls the expression of the GAL4 transcriptional activator in MuSCs, which in turn activates the expression of any gene placed under the control of UAS element. *Zfh1-Gal4* activity induces the expression of the *UAS-Flipase* and *UAS-RFP*. The Flipase enzyme recognizes FRT sites and removes the STOP cassette, allowing the expression of EGFP driven by the Ubi-p63E promoter in MuSCs and their progenies (Figure 1A). At low temperatures (18°C), tubGal80^ts^ inhibits Gal4 activity, whilst at high temperatures (29°C) the tubGal80^ts^ protein becomes non-functional. Thus, utilizing *tubGal80^ts^* to repress early Gal4 activity during development enables us to monitor MuSC activity in adult skeletal muscles through the comparison of Zfh1-Gal4 activity (RFP) and lineage-traced GAL4 activity (GFP), while avoiding potential early developmental issues arising from the improper modulation of target gene activities (Figure 1A).

**Figure 1.**
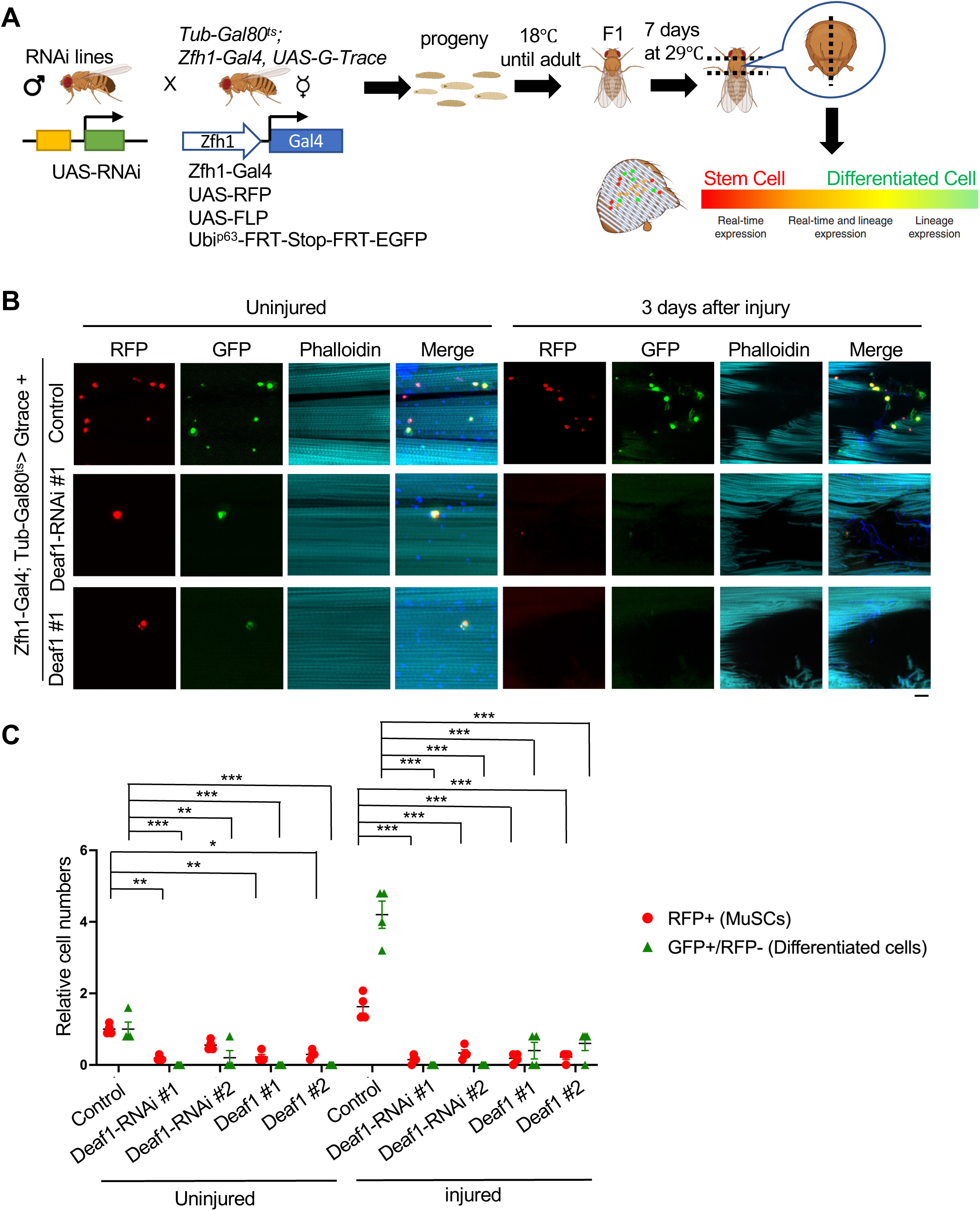
*Drosophila* Deaf1 regulates muscle regeneration. (A) Flowchart of the genetic screen. Crosses were set up between *Tub-Gal80^ts^; Zfh1-Gal4, UAS-G-Trace [72]* and different *UAS-RNAi* flies. Progenies were reared at 18 °C to avoid unintended RNAi expression during fly development. Adult flies were shifted to 29 °C for 7 days to allow *Zfh1-Gal4* expression that drives the expression of *RNAi*, *RFP*, and FLP recombinase specifically in MuSCs. MuSCs expressing FLP recombinase then excise the FRT-flanked stop cassette located between the Ubi-p63E promoter and EGFP. EGFP expression is heritably maintained in all daughter cells. Thus, RFP labels MuSCs expressing *Zfh1-Gal4* while GFP marks cells that are descendants of MuSCs. (B-C) Overexpression of *Deaf1* or *Deaf1-RNAi* leads to muscle regeneration defects. Representative images of *Drosophila* muscles at day 7 after G-trace was induced. In injured flies, muscles were stabbed with needles. Subsequently, muscles with or without injuries were stained by phalloidin (cyan) and DAPI (blue). MuSCs labeled by RFP (implying real-time Zfh1 expression) and differentiated cells labeled by GFP+/RFP- (implying lineal origin from a Zfh1-positive cell) were quantified in (C). Deaf1-RNAi #1 and #2 represent the independent shRNA lines with different targeting sequences. Both targeted Deaf1 and induced same phenotypes, suggesting that the phenotypes we observed were not due to off-target effects. For Deaf1 transgene #1 and #2, they are different constructs with or without tags. Both induced the same phenotypes, suggesting that the effects are not due to construct or plasmid insertion site issues. Scale bar: 10 µm.

One of the top candidates identified from the G-Trace screen was Deformed epidermal autoregulatory factor-1 (Deaf1). Deaf1 is an evolutionarily conserved transcription factor associated with autoimmune and neurological disorders [31, 32]. Two RNAi lines against *Deaf1* significantly decreased the numbers of MuSCs (RFP+ cells) as well as differentiated cells (GFP+/RFP- cells) under normal conditions compared to control (Figure 1B-C). In response to muscle injury, both MuSCs and differentiated cells were increased in the control flies, confirming that Zfh1-labeled MuSCs possess regeneration functions (Figure 1B-C). However, we did not observe this phenomenon in *Deaf1-RNAi*-expressing flies. The numbers of MuSC expressing *Deaf1-RNAi* and its lineage cells failed to increase in response to muscle injury (Figure 1B-C), suggesting that Deaf1 is required for the maintenance of MuSC pool and regeneration.

To further test the functions of *Deaf1* in muscle regeneration, we overexpressed *Deaf1* in MuSCs and expected that *Deaf1* overexpression may induce opposite effects from that of *Deaf1-RNAi*. Unexpectedly, we found that *Deaf1* overexpression in MuSCs dramatically diminished both RFP+ and GFP+/RFP- cells (Figure 1B-C), suggesting that *Deaf1* overexpression disrupts muscle regeneration. During muscle injury, MuSCs overexpressing *Deaf1* did not proliferate and differentiate as compared to injured control (Figure 1B-C). Thus, these results suggest that *Deaf1* depletion and its overexpression both resulted in regeneration defects. To further investigate the Deaf1-induced MuSC defects, we induced G-trace labeling in MuSCs expressing *Deaf1* or *Deaf1-RNAi*, and quantified MuSCs (RFP+ cells) and differentiated cells (GFP+/RFP- cells) at successive intervals after induction (Figure S1). We found that, in contrast to control, the numbers of MuSCs and differentiated cells were both reduced by expression of *Deaf1* or *Deaf1-RNAi* within 7 days of induction (Figure S1). Together, these results suggest that alteration of *Deaf1* expression leads to cell elimination, highlighting that the fine-tuning of Deaf1 is critical for the maintenance of MuSC pool and determining cell fate of MuSCs.

### Autophagy is critical for Deaf1-regulated MuSC homeostasis

The critical roles of autophagy in MuSCs have been addressed by several independent studies. Some studies suggest that autophagy is crucial for maintaining stemness, and low levels of autophagy has been observed in aging MuSCs [13, 33]. However, hyperactive autophagy has been shown to reduce the proliferative capacity of MuSCs, which plays an important role in the early regeneration of damaged skeletal muscle in myotonic dystrophy type 1 (DM1) [34]. Therefore, these findings suggest that both deficient and excessive autophagy in MuSCs result in a pathological cascade and lead to muscle atrophy symptoms [3, 35].

Decreased and accelerated autophagy in MuSCs leads to similar defects caused by *Deaf1* alteration, raising the possibility that Deaf1 may be involved in the regulation of autophagy. In our previous genetic screen, we identified that *Deaf1* genetically interacts with *Atg1* and that depletion of *Deaf1* induces autophagy in larval fat body [18]. Therefore, we investigated if Deaf1 may also regulate autophagy in MuSCs. *Drosophila* Atg8a protein, an orthologue of mammalian LC3, is an autophagosomal reporter and was expressed in MuSCs with or without chloroquine (CQ) treatment. CQ inhibits autophagosome-lysosome fusion and thus increases mCherry-ATG8a puncta under normal conditions [36] (Figure 2A-B). *Deaf1-RNAi* expression increased mCherry-ATG8a puncta with or without CQ treatment. In contrast, MuSCs overexpressing *Deaf1* displayed decreased levels of ATG8a puncta (Figure 2A-B), suggesting that Deaf1 is a negative regulator of autophagy. Consistently, our GFP-mCherry-Atg8a reporter assay also revealed increased levels of both autophagosomes (mCherry+; GFP+ vesicles) and autolysosomes (mCherry+; GFP-vesicles) in *Deaf1-knockdown* cells, but decreased levels in both autophagosomes and autolysosomes in *Deaf1*-overexpressing MuSCs (Figure S2), suggesting that *Deaf1* depletion induces autophagy while the overexpression of *Deaf1* suppresses autophagy.

**Figure 2.**
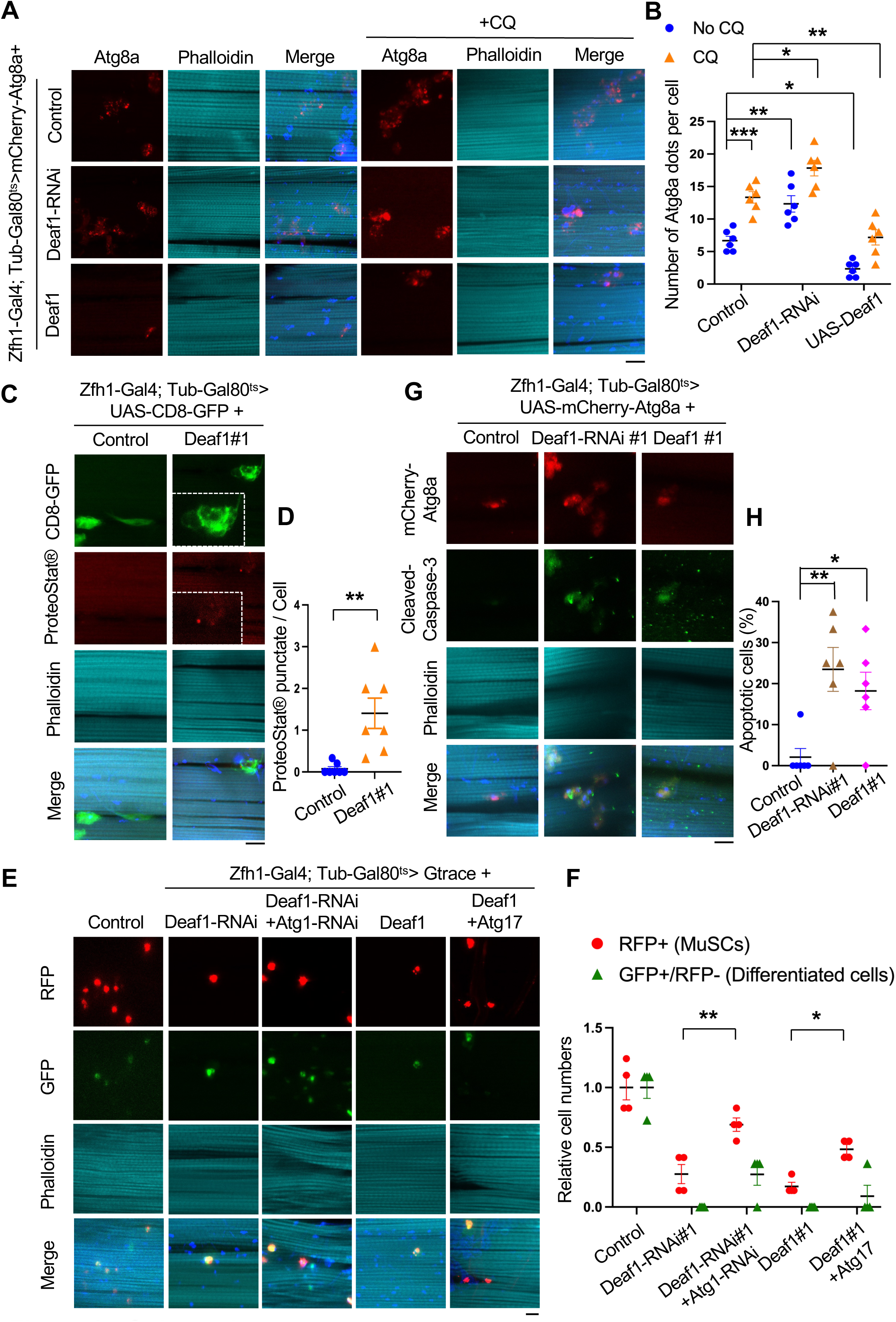
Deaf1-regulated autophagy controls MuSC maintenance and differentiation in *Drosophila*. (A-B) *Drosophila* Deaf1 negatively regulates autophagy. *Drosophila* adult muscles from *tubGal80^ts^; Zfh1-Gal4, UAS-mCherry-Atg8a* controls or flies expressing *Deaf1* or *Deaf1-RNAi* fed with or without 10 mM Chloroquine for three days were stained with phalloidin (cyan) and DAPI (blue). Atg8a punctate were quantified in (B). Compared to control, *Deaf1-RNAi* increased Atg8a punctate numbers while Deaf1 overexpression suppressed Atg8a-labeled autophagosome formation. Scale bar: 10 µm. (C-D) Deaf1 overexpression increases protein aggregates. Adult muscles from *tubGal80^ts^; Zfh1-Gal4, UAS-CD8-GFP* controls or flies expressing *Deaf1* were stained with ProteoStat® aggresome dye (red) which detects protein aggregates, phalloidin (cyan), and DAPI (blue) (C). ProteoStat® signals were quantified in (D). Scale bar: 10 µm. (E-F) Modulation of autophagic activity restores *Deaf1* overexpression*-* or *Deaf1-RNAi*-induced muscle regeneration defects. Adult muscles from flies expressing indicated transgenes were stained with phalloidin (cyan) and DAPI (blue) (E). Scale bar: 10 µm. MuSCs labeled by RFP and differentiated cells labeled by GFP+/RFP- were quantified in (F). (G-H) Changes of *Deaf1* expression levels trigger MuSC deaths. Adult muscles from flies as in (A) were stained with anti-cleaved Caspase-3 (green), phalloidin (cyan), and DAPI (blue) (G). Apoptotic cells labeled by cleaved-caspase-3 were quantified in (H). Scale bar: 10 µm.

Autophagy is required for maintaining proteostasis while a high level of autophagy induces cell death [13, 37, 38]. To explore the functions of autophagy in *Deaf1*-regulated MuSCs, we next examined whether *Deaf1* expression leads to proteotoxicity and if changes in Deaf1 levels results in apoptosis. First, we stained the cells with PROTEOSTAT^®^ aggregation dye to monitor the protein aggregates in MuSCs [39]. As shown in Figure 2C, increased protein aggregates were observed in *Deaf1*-overexpressing MuSCs (Figure 2C-D). Atg1 and Atg17 are both key components of Atg1 complex. Knockdown of *Atg1* represses autophagy while overexpression of *Atg17* enhances it [40, 41]. As Deaf1 suppressed autophagy, we co-expressed Atg17 to enhance autophagy and tested if Atg17 could rescue Deaf1-induced effects. Indeed, expression of *Atg17* partially rescued *Deaf1*-induced MuSC defects (Figure 2E-F). Suppression of autophagy by co-expression of *Atg1-RNAi* increased *Deaf1*-depleted MuSCs numbers and restored their differentiation activity, as compared to *Deaf1-RNAi* expression alone (Figure 2E-F). These results suggest that autophagy mediates downstream functions of Deaf1. In addition, apoptotic cells were observed in MuSCs expressing *Deaf1* or *Deaf1-RNAi* (Figure 2G-H). Together, these results demonstrate that autophagy functions downstream of Deaf1 and that the Deaf1-autophagy signaling axis plays critical roles in regulating MuSCs.

### Mammalian Deaf1 regulates autophagy in both C2C12 myoblasts and primary MuSCs

We next tested if the roles of Deaf1 in regulating autophagy are conserved in mammals. We generated C2C12 myoblasts stably expressing *Deaf1* and found that *Deaf1* expression suppressed autophagosome formation in the presence or absence of the lysosomal inhibitor Bafilomycin A1 (BafA1) (Figure 3A-B). Consistent with this result, *Deaf1*-expressing C2C12 showed an increase in protein aggregates, as assessed by PROTEOSTAT^®^ aggresome dye as well as by anti-Ubiquitin antibody (Figure 3C-D and S3A), suggesting that Deaf1 is a negative regulator of autophagy required for maintaining proteostasis. We further observed that C2C12 myoblasts expressing *Deaf1* failed to differentiate into myocytes, as indicated by immunostaining of Myosin heavy chain (MHC), a marker of myocytes (Figure 3E-F). Accordingly, the fusion index showed that significantly lower myocyte fusion occurred in C2C12 myoblasts expressing *Deaf1*, as compared to control (Figure 3G). *Deaf1*-expressing C2C12 cells also exhibited decreases in proliferation rate (Figures 3H). To further confirm these results, we isolated primary MuSCs from mouse muscles and expressed *Deaf1* in these cells. *Deaf1* expression in primary MuSCs reduced LC3 punctate formation, demonstrating that Deaf1 suppresses autophagy (Figure 3I-J). Consistent with C2C12 myoblast results, primary MuSCs expressing *Deaf1* also showed differentiation defects (Figure 3K-M) as well as increased protein aggregates (Figure 3N-O and S3B). Taken together, these results demonstrate that Deaf1 negatively regulates autophagy and is critical for proliferation and differentiation of MuSCs.

**Figure 3.**
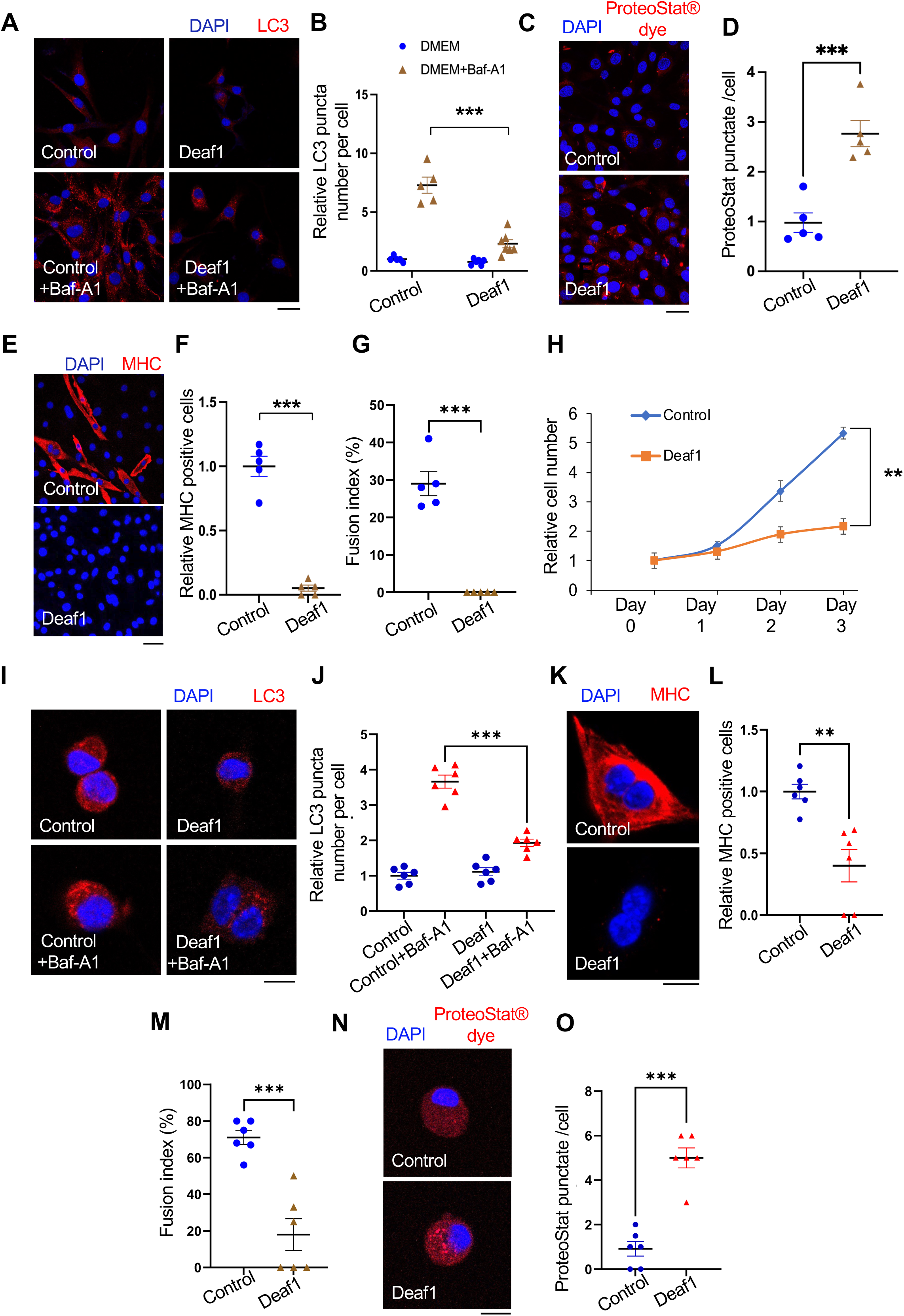
Deaf1 overexpression suppresses autophagy and inhibits MuSC proliferation and differentiation in mammals. (A-B) Deaf1 inhibits autophagosome formation in C2C12 myoblasts. C2C12 cells stably infected with lentivirus expressing *Deaf1* were cultured in DMEM with or without Bafilomycin A1 (BafA1) treatment for 4 hrs and then subjected to immunofluorescence with anti-LC3B (red) and DAPI (blue) (A). LC3 puncta were quantified in (B). Scale bar: 20 µm. (C-G) Deaf1 overexpression induces protein aggregate formation and blocks myoblast differentiation. C2C12 myoblasts stably infected with lentivirus expressing *Deaf1* were cultured in DMEM with 10% FBS or with 2% horse serum (for induction of muscle differentiation) and subjected to immunofluorescence with ProteoStat® aggresome dye (C, red) to monitor protein aggregates or anti-MHC antibody (E, red) to indicate differentiated cells. Nuclei were stained with DAPI (blue). Protein aggregates and differentiated cells were quantified in (D) and (F) respectively. Fusion index which is calculated as the percentage of nuclei incorporated in the myotubes relative to the total number of nuclei was shown in (G). Scale bar: 20 µm (C) and 50 µm (E). (H) *Deaf1* expression represses cell growth. Cell counts on day1-3 after seeding. (I-J) *Deaf1* decreases autophagy flux in isolated MuSCs from mouse muscles. Purified MuSCs infected with lentivirus expressing *Deaf1* were treated with or without Baf-A1 for 4hrs and subjected to immunofluorescence with anti-LC3B (red) and DAPI (blue) (I). LC3 puncta were quantified in (J). Scale bar: 10 µm. (K-O) Deaf1 represses MuSC differentiation as well as increases protein aggregates. Purified MuSCs infected with lentivirus expressing *Deaf1* were cultured with or without 2% horse serum to induce muscle differentiation and were subjected to immunofluorescence with anti-MHC antibody (K, red) or ProteoStat® aggresome dye (N, red). Nuclei were stained with DAPI (blue). Myocytes, fusion index, and protein aggregates were quantified in (L), (M) and (O) separately. Scale bar: 10 µm.

To further verify Deaf1 functions using a loss of function approach, we generated *Deaf1* Knockout (KO) C2C12 cells (Deaf1 KO #1 and #2). In contrast to *Deaf1* overexpression, we observed significant increases in LC3 punctate numbers in *Deaf1-KO* C2C12 myoblasts when cells were treated with Baf-A1 (Figure 4A-B). Furthermore, both clones of *Deaf1KO* cells showed remarkedly reduced proliferation assessed by cell growth over three days (Figure 4C). *Deaf1KO* cells also exhibited reduced protein aggregates indicated by PROTEOSTAT^®^ staining as well as defects in differentiation assessed by immunostaining of Myosin heavy chain and myocyte fusion index assay (Figure 4D-F). Consistent with *Drosophila* results, suppression of autophagy by expressing shAtg7 in *Deaf1KO* cells partially reversed the *Deaf1KO*-induced effects (Figure 4D-F). These results suggest that *Deaf1* deficiency induces autophagosome formation and results in MuSC proliferation and differentiation defects. Similarly, knockdown of *Deaf1* in primary MuSCs purified from mouse muscles exhibited increased LC3 punctate when MuSCs were treated with Baf-A1 (Figure 4G-H). Consistently, *Deaf1* depletion also led to a decrease in protein aggregates as well as differentiation defects (Figure 4I-K). Autophagy inhibition by expressing shAtg7 was able to rescue shDeaf1-induced effects (Figure 4G-K). Together, our findings demonstrate that *Deaf1* loss of function enhances autophagy, leading to defects in MuSC proliferation and differentiation.

**Figure 4.**
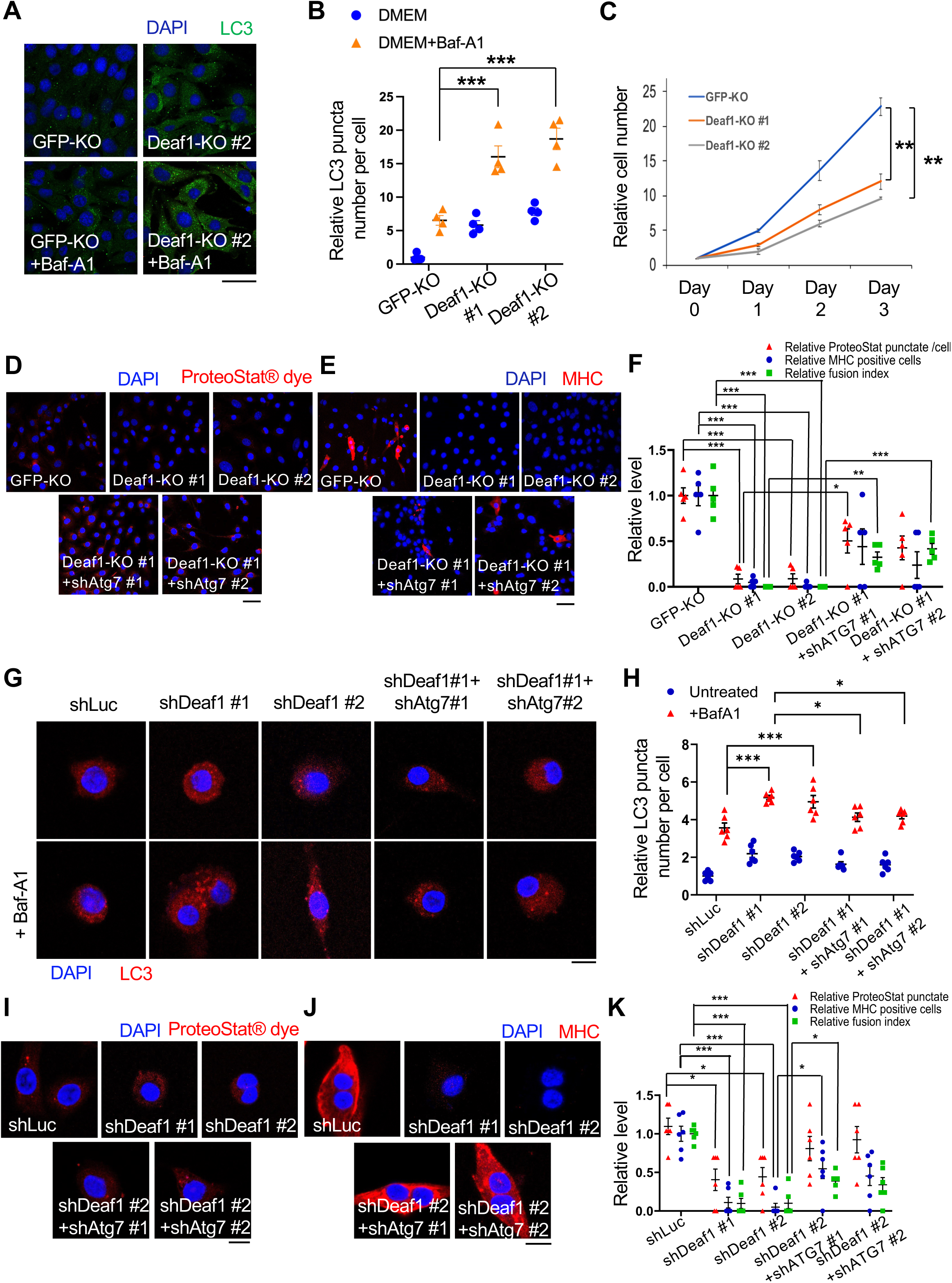
*Deaf1* depletion induces autophagy and represses proliferation and differentiation of C2C12 myoblasts and primary MuSCs. (A-B) *Deaf1* deficient myoblasts exhibit increased autophagy. GFP- or *Deaf1*-knockout clones of C2C12 cells were cultured in DMEM with or without Bafilomycin A1 (BafA1) for 4hrs and then subjected to immunofluorescence with anti-LC3B antibody(green) and DAPI (blue) (A). LC3 puncta were quantified in (B). Scale bar, 20 µm. (C) *Deaf1* depletion suppresses cell growth. Cell counts on day1-3 after seeding. (D-F) *Deaf1*-knockout C2C12 myoblasts exhibit reduced protein aggregates and defects in differentiation due to autophagy overactivation. *Deaf1*-*KO* C2C12 infected with or without lentivirus expressing *shAtg7* cultured in DMEM with 10% FBS or with 2% horse serum (for induction of muscle differentiation) and subjected to immunofluorescence with ProteoStat® aggresome dye (D, red) to monitor protein aggregates or anti-MHC antibody (E, red) to indicate differentiated cells. Protein aggregates, differentiated cells, and fusion index, were quantified in (F). Nuclei were stained with DAPI (blue). Scale bar: 50 µm. (G-H) *Deaf1* knockdown increases autophagy flux in MuSCs. Purified MuSCs from mouse muscles infected with lentivirus expressing *shLuc* (control), *shDeaf1, or shDeaf1* together *with shAtg7,* were treated with or without Baf-A1 for 4hrs and subjected to immunofluorescence with anti-LC3B antibody (red) and DAPI (blue) (G). LC3 puncta were quantified in (H). Scale bar: 10 µm. (I-K) *Deaf1* knockdown decreases protein aggregates and blocks MuSC differentiation. Purified MuSCs infected with lentivirus expressing s*hDeaf1* were cultured with or without 2% horse serum, subjected to immunofluorescence with ProteoStat® aggresome dye (I, red) or anti-MHC antibody (J, red), and quantified in (K). Nuclei were stained with DAPI (blue). Scale bar: 10 µm.

### Deaf1 binds to promoters of autophagy-related genes and suppresses their expression

To address how Deaf1 regulates autophagy, we subsequently conducted a chromatin immunoprecipitation assay with DNA sequencing (ChIP-seq) analysis in C2C12 myoblasts to identify the potential downstream targets of Deaf1. This analysis led to the identification of 608 Deaf1 binding events in C2C12 cells (IDR > 5%) (Table S1). *De novo* motif analysis identified the presence of a motif that is consistent with the known Deaf1 motif, confirming the quality of our data [31, 42] (Figure. 5A). The clear majority (96%) of these events were found at gene promoters (Figure 5B), consistent with the role of Deaf1 in transcriptional regulation.

**Figure 5.**
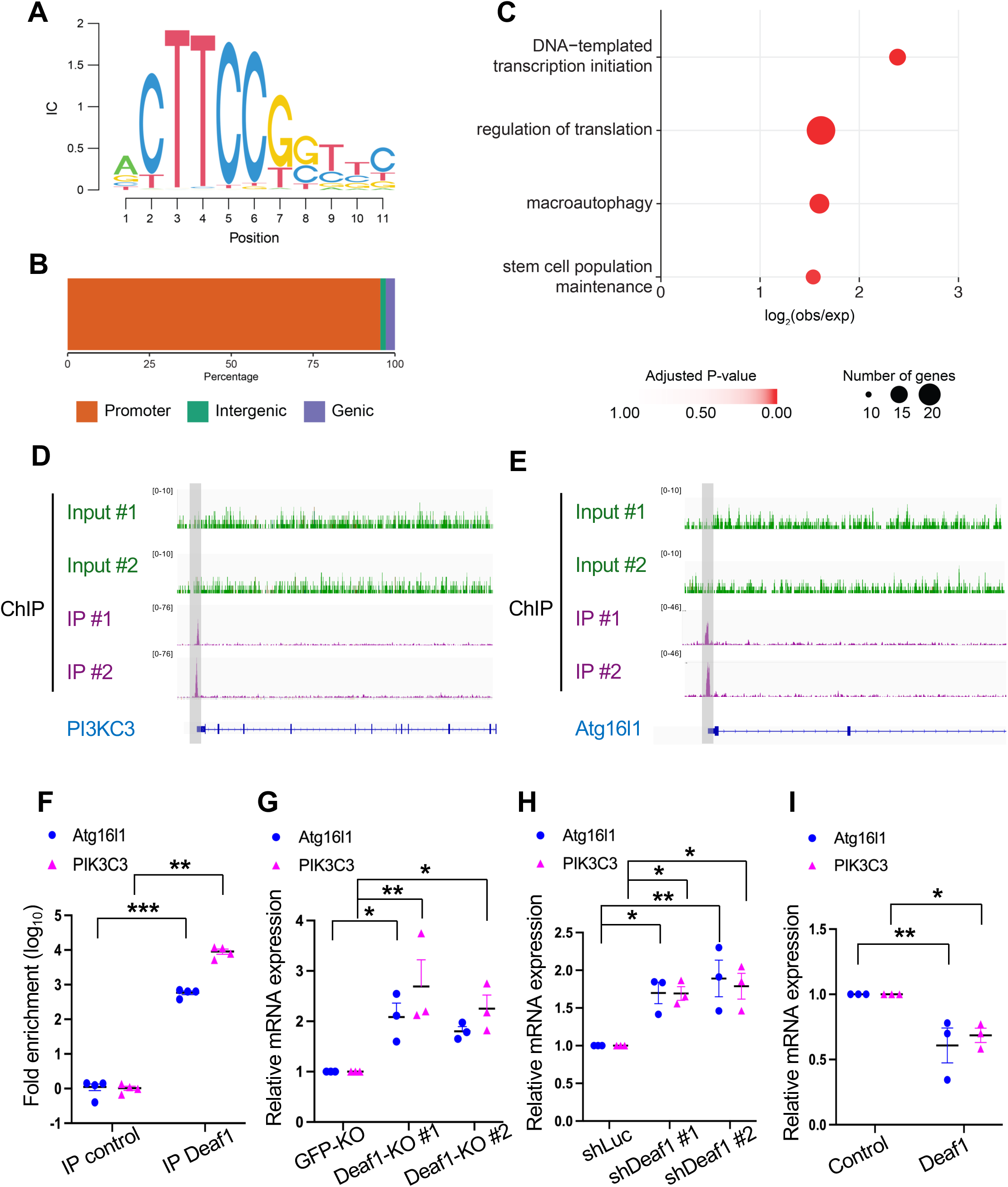
Deaf1 binds to promoter regions of *Atg16L1* and *PIK3C3* genes and suppress their transcription. (A) Consensus binding motif sequence identified from ChIP-seq data using anti-Deaf1 antibody. (B) Distribution of Deaf1 binding sites from ChIP-seq. (C) Gene ontology enrichment analysis of ChIP-seq data. (D-E) Example browser images for PI3KC3 (D) and Atg16l1 (E) from ChIP-seq in C2C12 myoblast cells using anti-Deaf1 antibody - data from two independent experiments shown. (F) Deaf1 occupancy at promoter regions of *PI3KC3* and *Atg16l1* genes was revealed by ChIP-qPCR. (G-I) Deaf1 suppresses *PI3KC3 and Atg16l1* expression. Increased expression of *PI3KC3 and Atg16l1* transcripts were observed in *Deaf1*-knockout cells (G) or cells expressing *shDeaf1* (H), but decreased expressions occurred in cells stably expressing *Deaf1* (I).

Interrogation of the genes bound by Deaf1 identified significant enrichments for core transcriptional and translational processes, as well as genes involved in stem cell maintenance and autophagy (Figure 5C) (Table S2). Amongst autophagy-related genes (Atg), we identified two Atg genes, *Atg16l1* and *PI3KC3*, as Deaf1 downstream targets. Our ChIP-seq analysis identified the presence of Deaf1 peaks at the promoter region of *PI3KC3* and *Atg16l1* (Figure 5D-E). The ChIP-qPCR results further confirmed Deaf1’s binding to the promoter regions of *Atg16l1* and *PIK3C3*, with a significant increase in fold enrichment in comparison to the control (Figure 5F), suggesting that Deaf1 can bind to the promoter regions of Atg16l1 and PI3KC3 genes.

Next, we explored the effects of Deaf1 on *Atg16l1* and *PI3KC3* mRNA expression. *Atg16l1* and *PI3KC3* mRNA levels were significantly increased in *Deaf1*-knockout and -knockdown cells (Figure 5G-H) but decreased in *Deaf1*-expressing cells (Figure 5I). Thus, our results suggest that Deaf1 can inhibit the expression of key autophagy regulators, Atg16l1 and PI3KC3, by targeting their promoters.

### Deaf1 acts downstream of FOXOs and mediates FOXO-dependent autophagy

A recent ChIP-seq analysis identified Deaf1 as a potential target of FOXO in *Drosophila* muscle [43]. More interestingly, the promoter of Deaf1 is bound by FOXO in young flies, but not in old flies, suggesting that Deaf1 may be a downstream target of FOXO and that FOXO binding to Deaf1 promoter may be inhibited upon muscle aging [43].

To determine whether FOXOs are involved in Deaf1-regulated autophagy, we examined the effects of *Deaf1* overexpression or *Deaf1* knockdown on FOXO-dependent autophagy. C2C12 myoblasts treated with LOM612, a FOXOs activator [44], along with Baf-A1, showed increased LC3 punctate numbers compared to cells treated with Baf-A1 alone (Figure S4A-B). *Deaf1* overexpression was sufficient to significantly reduce LOM612-increased LC3 punctate formation (Figure S4A-B), suggesting that *Deaf1* expression can suppress FOXO activation-induced autophagy. In contrast, the treatment of C2C12 cells with AS1842856, a FOXO1 inhibitor [45], reduced LC3 punctate numbers (Figure S4A-B). We used *Deaf1* shRNA to downregulate *Deaf1* and found that decreased *Deaf1* expression can reverse the AS1842856-induced autophagy defects, showing that Deaf1 is required for the FOXO inhibitor-induced effects (Figure S4A-B). These results suggest that that Deaf1 functions downstream of FOXOs and mediates FOXOs-dependent autophagy.

### FOXOs binds to the promoter of *Deaf1* and repress its transcription

FOXO transcription factors bind specific double-stranded DNA sites including the DAF-16-binding element (DBE, 5′-GTAAA(T/C)AA-3′) and the insulin responsive element (IRE, 5′-(C/A)(A/C)AAA(C/T)AA-3′) [46]. Intriguingly, we identified an IRE at the promoter of *Deaf1* in both *Drosophila* and mouse (Figure 6A). Our ChIP-qPCR results further revealed that *Drosophila* FOXO can bind to *Deaf1* promoter in young muscle, but not in old muscles (Figure 6B), consistent with a previous study [43]. In mouse C2C12 myoblasts, both FOXO1 and FOXO3 can bind to the *Deaf1* promoter (Figure 6C). Together, these results show that Deaf1 is a target of FOXOs.

**Figure 6.**
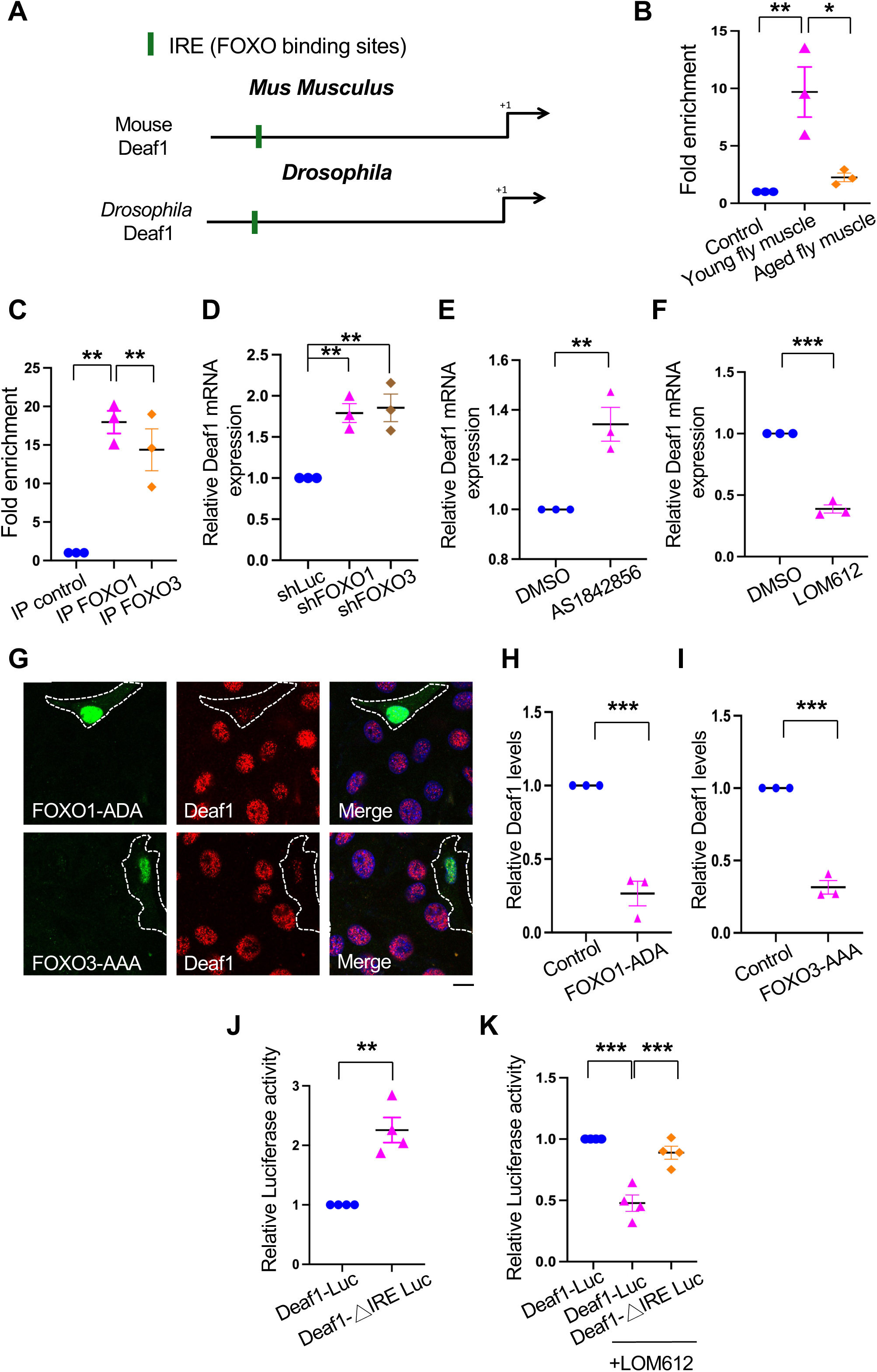
FOXOs target and repress Deaf1 expression. (A) A putative FOXO binding site (green box; Insulin Responsive Element [IRE]) located in the proximal (2 kb) promoter region of fly and mouse *Deaf1*. Arrows denote ATG. (B-C) FOXO transcription factors bind to the *Deaf1* promoter. ChIP was performed using young and old fly thorax (ages: 2 vs. 5 weeks) (B) or using C2C12 cells (C), followed by qPCR assay. (D-F) FOXOs suppress *Deaf1* mRNA expressions in C2C12 myoblasts. RNAs from C2C12 myoblasts stably infected with lentivirus expressing *shLuc, shFOXO1,* or *shFOXO3*, or treated with AS-1842856 (a FOXO1 inhibitor) or LOM612 (a FOXOs activator) were subjected to qPCR assays. (G-I) Expression of constitutively active FOXO1 or FOXO3 decreases Deaf1 protein levels. C2C12 myoblasts were transfected with activated FOXO1 (FOXO1-ADA) or FOXO3 (FOXO3-AAA) (green) were subjected to immunofluorescence with anti-Deaf1 antibody (red) and DAPI (blue) (G). Deaf1 signals in control C2C12 cells or cells expressing active FOXO1 (H) or FOXO3 (I) were quantified. Scale bar: 10 µm. (J-K) Luciferase assays using wild-type (wt) and mutant versions (ΔIRE) of the Deaf1 promoter. Removal of FOXOs binding sites (ΔIRE mutant reporters) increases the transcriptional activity of Luciferase reporters (J). The LOM612 treatment (a FOXOs activator) suppresses the *Deaf1* transcription which is reversed by the removal of FOXO binding sites (K).

To determine the effects of FOXOs on *Deaf1*, we next used two independent methods to modulate FOXO activity and assessed the transcriptional changes of *Deaf1*. C2C12 myoblasts were transfected with *FOXO1* or *FOXO3* shRNAs or treated with AS1842856, a FOXO1 inhibitor. We found that depletion or inactivation of FOXOs significantly increased *Deaf1* mRNA levels, suggesting FOXOs target and inhibit *Deaf1* expression (Figure 6D-E). In contrast, C2C12 cells treated with LOM612, a FOXOs activator, exhibited decreased *Deaf1* mRNA levels, as compared to control (Figure 6F). To further confirm these results, we expressed a constitutively active form of either FOXO1 (FOXO1-ADA) or FOXO3 (FOXO3-AAA) in C2C12 myoblasts and found that Deaf1 protein level was strongly reduced, as assessed by immunofluorescence (Figure 6G-I). Together, these data demonstrate that FOXOs suppress Deaf1 expression by DNA binding to its promoter. Interestingly, we observed the same phenomenon in C2C12 myocytes, LL2 (lung), and HCT116 (colon) cells when cells were treated with AS1842856 or LOM612 (Figure S4C). However, no significant changes in *Deaf1* levels were detected in HEK293T (kidney) cells (Figure S4C). These results suggest that the FOXO-Deaf1 axis also exists in other tissues but may not so in the kidney.

To further confirm whether FOXOs directly suppresses *Deaf1* transcription, we cloned the mouse *Deaf1* promoter regions with or without deletion of FOXO binding sites (Deaf1-△IRE) and linked them to Luciferase. These Luciferase transcriptional reporter assays revealed that deletion of FOXO binding sites in mouse *Deaf1* promoter regions increased luciferase activities (Figure 6J) and that LOM612 treatment reduced Deaf1 transcription, an effect that was reversed by the deletion of FOXO binding sites (Figure 6K). Thus, these findings demonstrate that FOXO directly binds to the promoter regions of Deaf1 and inhibits its transcription. Together, our results identify a novel transcriptional cascade, FOXOs-Deaf1-autophagy which regulate muscle regeneration.

### Changes in *Deaf1* expression during aging and cancer cachexia

MuSCs dysfunction has been linked to multiple human muscle diseases, including sarcopenia and cancer cachexia [3]. We thus tested the roles of Deaf1 in these diseases by examining *Deaf1* expression levels. Intriguingly, we observed substantial increases in *Deaf1* mRNAs in muscles from old flies (ages: 60 days) in comparison to that of young flies (ages: 7 days) (Figure 7A). We next purified MuSCs from young (ages: ∼3 months) and old (ages:>18 months) mice respectively and found that *Deaf1* mRNA levels were higher in old MuSCs than that in young MuSCs, suggesting that *Deaf1* is increased in aging MuSCs (Figure 7B). Consistent with our previous result, we also found that mRNAs of FOXO1 and FOXO3 were reduced in aging MuSCs (Figure S5A-B). We subsequently examined previously published RNA-seq datasets to look for evidence of changes in *Deaf1* expression in sarcopenic patients (Figure S5C). In both Jamaican and Singaporean patients classified as having sarcopenia, *Deaf1* expressions were slightly upregulated, although these were not significant (Figure S5C). These may be due to the presence of *Deaf1* mRNA in the muscles which partially masks its changes in MuSCs.

**Figure 7.**
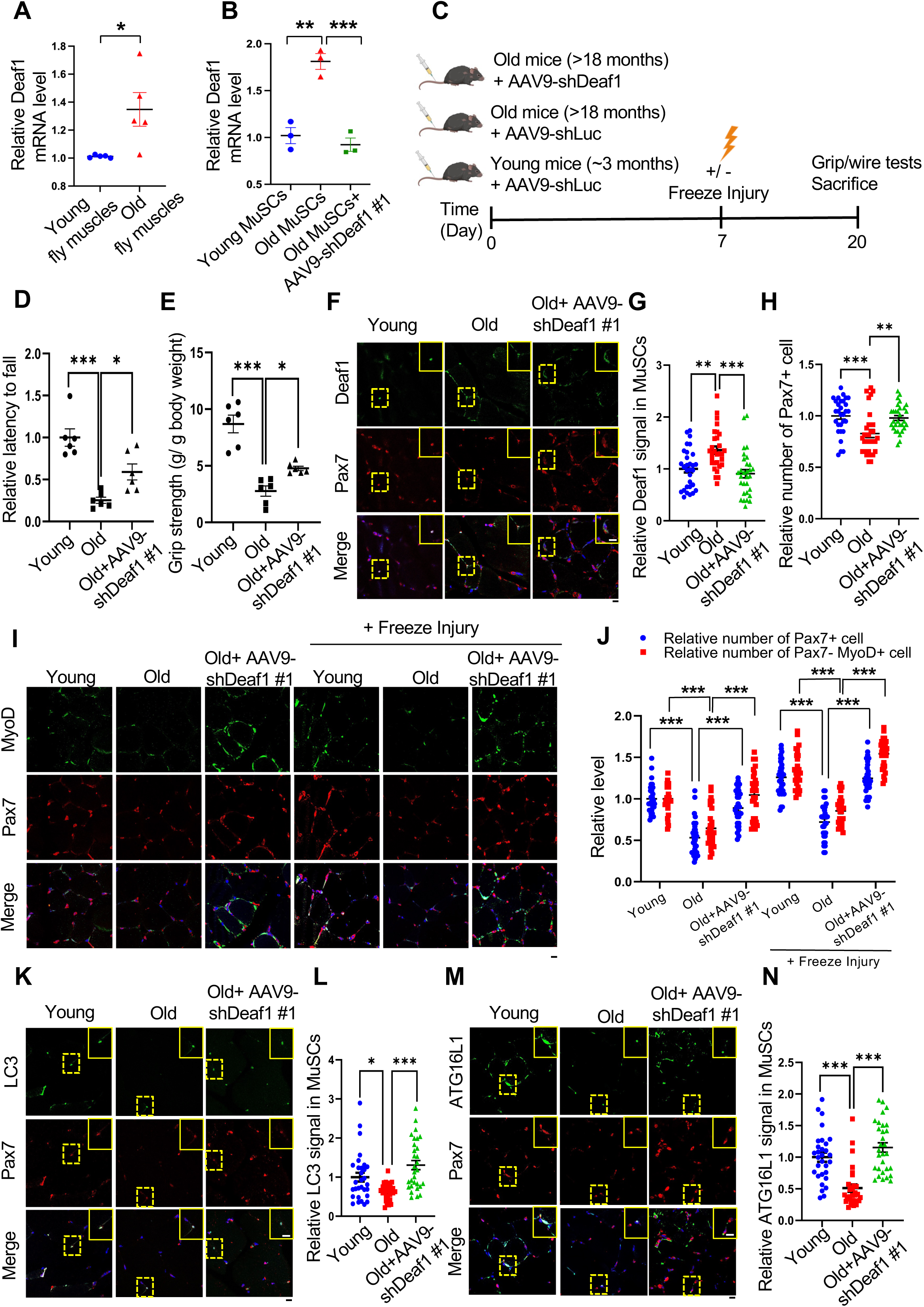
Modulation of Deaf1 expression levels restores muscle functions in old muscles. (A-B) *Deaf1* mRNA is increased in aging MuSCs. RNA extracts from thorax of young (age: 1 week) or old (age: 6 weeks) flies (A) as well as isolated MuSCs from young (age: ∼3 months) or old (age: 18 months) soleus muscles with or without AAV9-shDeaf1 injection (B) were subjected to qPCR analysis. (C) Flow chart showing the schema of mouse experiment using AAV9. (D-E) Expression of shDeaf1 improves muscle function during aging. Young (∼3 months) or aged (>18 months) C57BL/6J mice were intravenously injected with AAV-shLuc (control) or AAV9-shDeaf1 and subjected to the wire hanging (D) and grip strength (E) tests on day 20. (F-L) Reduction of *Deaf1* expression increases autophagy and MuSC regeneration in aging muscles. Immunostaining of soleus muscles from young or aged mice with or without freeze injury using anti-Deaf1 (green) (F), anti-MyoD (green) (I), anti-LC3 (green) (K), anti-ATG16L1 (green) (M), and anti-Pax7 (red) antibodies on day 20 post AAV9-mediated transduction. Deaf1 (G), Pax7 (H and J), Pax7-/MyoD+ (J), LC3 (L), and ATG16L1 (N) signals in MuSCs were quantified. Scale bar: 10 µm.

In contrast, our qPCR data showed that *Deaf1* mRNAs were decreased in cachectic muscles from flies bearing *yki^3SA^*-gut tumors [47, 48] (Figure 9A). In a mouse cachexia model where LL2 lung cancer cells were implanted subcutaneously and induced muscle wasting, lower *Deaf1* expression levels were detected in purified MuSCs from mice with LL2 tumors compared to control mice (Figure 9B). Consistently, both FOXO1 and FOXO3 mRNAs were increased in cachectic MuSCs (Figure S5D-E). Furthermore, in human cachectic muscles, *Deaf1* expression was significantly decreased (Figure S5F), suggesting that *Deaf1* expression is suppressed during cancer cachexia. Together, these results from different model systems revealed that Deaf1 is upregulated in sarcopenia and downregulated in cancer cachexia, indicating that mechanisms underlying these muscle diseases are different, even though both show muscle wasting and a reduction in MuSC numbers.

### Modulation of Deaf1 promotes muscle regeneration under aging and cancer cachectic conditions

To determine whether reduction of *Deaf1* levels can improve muscle functions, we used adeno-associated virus serotype 9 (AAV9) to transduce MuSCs [49] or intramuscular injection of FOXO activator/inhibitor into hindlimb muscles respectively to alter *Deaf1* levels in MuSCs. Firstly, we measured the muscle strengths and functions of young (age: ∼3 months) and old mice (age: >18 months) using grip strength assay and wire hanging test (Figure 7C-E). The grip strength assay is a widely used method to measure maximal muscle strength in rodents [50], while skeletal muscle functions (endurance and muscle tone) is assessed by a wire hanging test [51]. Compared to young mice, aged mice showed reduced grip strength and wire holding ability (Figure 7D-E). Significantly, AAV9-mediated delivery of *shDeaf1* through intravenous (IV) injection into aged C57BL/6J mice decreased *Deaf1* expression in MuSCs and reversed the aging-reduced muscle functions and strengths (Figure 7B, D, and E), suggesting that *Deaf1* knockdown can rescue aging-induced muscle wasting. Our immunofluorescence staining of Deaf1 in soleus muscles further revealed that Deaf1 protein was highly enriched in MuSCs labeled by Pax7 in young muscles (Figure 7F-G). In old muscles, Deaf1 levels in MuSCs were increased, MuSCs numbers were reduced, and differentiated myoblasts labeled by Pax7-/MyoD+ were decreased (Figure 7F-J). AAV9-mediated *Deaf1* knockdown not only reduced the Deaf1 levels in MuSCs but also increased MuSC numbers and differentiated myoblasts (Pax7-/MyoD+ cells) with or without freeze-injury (Figure 7F-J), suggesting that *Deaf1* depletion promotes muscle regeneration during aging. Furthermore, compared to young MuSCs, LC3 puncta and ATG16L1 levels were reduced in aged MuSCs, a phenomenon reversed by expression of *Deaf1* shRNA (Figure 7K-N). Consistently, PIK3C3 levels were also reduced in old muscles, but increased by shDeaf1 expression (Figure S6A). These results suggest that Deaf1-regulated autophagy controls MuSC activity upon aging.

To further confirm these results, we intramuscularly injected LOM612, a FOXOs activator, into the hindlimb muscle of old mice to decrease *Deaf1* expression in MuSCs (Figure 8A). LOM612-mediated Deaf1 reduction increased MuSCs numbers indicated by Pax7 staining, differentiated myoblasts labeled by Pax7-/MyoD+, and autophagy levels assessed by LC3 puncta (Figure 8B-H). Accordingly, we also observed increases in ATG16L1 and PIK3C3 following LOM612 treatment (Figure 8I-J and S6C-D). Together, these findings demonstrate that FOXO-Deaf1-autophagy signaling plays key roles in MuSCs dysregulation upon aging, and that suppression of Deaf1 can restore muscle regeneration and improve muscle functions during aging.

**Figure 8.**
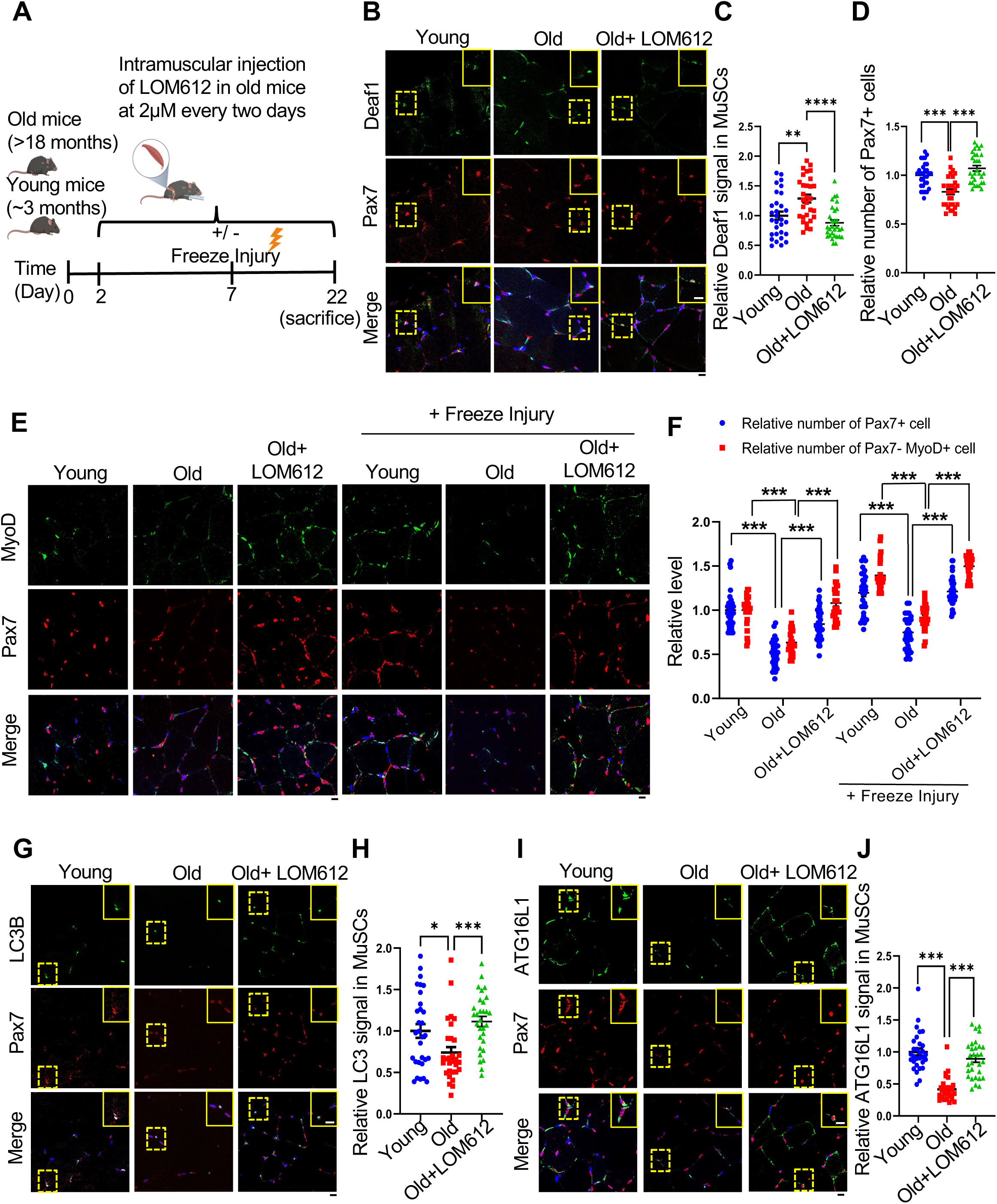
Deaf1 reduction induced by FOXO activation restores autophagy and improves muscle regeneration in aged MuSCs. (A) Schematic representation of the experimental design. (B-J) Young, aged mice, or aged mice with intramuscular injection of a FOXO activator, LOM612, were subjected to freeze injuries, followed by immunostaining using anti-Deaf1 (green) (B), anti-MyoD (green) (E), anti-LC3 (green) (G), anti-ATG16L1 (green) (I), and anti-Pax7 (red) antibodies. Deaf1 (C), Pax7 (D and F), Pax7-/MyoD+ (F), LC3 (H), and ATG16L1 (J) signals in MuSCs were quantified. Scale bar: 10 µm.

Next, we examined the roles of Deaf1 in cancer cachexia. We subcutaneously implanted mice with a cachectic lung cancer cell line LL2, to induce cancer-induced muscle wasting, and delivered AAV9-Deaf1 via intravenous (IV) injection (Figure 9B-C). AAV9-Deaf1 increased *Deaf1* level in MuSCs (Figure 9B-C), but did not affect LL2 tumor volume and weight, demonstrating the specificity of AAV9 (Figure S7A-B). Mice bearing LL2 cachectic tumors exhibited significantly reduced grip strength and wire holding ability (Figure 9D-E). This cancer-induced muscle atrophy was rescued by AAV9-Deaf1 (Figure 9D-E). Persistent expression of Pax7 and increased cell death have been observed during cancer cachexia [22, 23]. In soleus muscles from mice with LL2 tumors, we observed that Deaf1 expression in MuSCs was decreased and Pax7+ cells were increased (Figure 9F-H). However, differentiated cells labeled by Pax7-/MyoD+ were not increased and apoptosis were detected by Apoptag assay in cachectic muscles (Figure 9I-J). Moreover, LC3, ATG16L1, and PIK3C3 expressions were enhanced under cachexia condition, suggesting that cachectic MuSCs showed high levels of autophagy and exhibited defects in muscle differentiation during cancer cachexia (Figure 9K-N and S7C-D). Importantly, these cachectic cancer-induced effects were reversed by AAV9-mediated *Deaf1* expression (Figure 9F-N and S7C-D). We further verified these results by intramuscular injection of AS1842856, a FOXO1 inhibitor (Figure 10A). Consistent with our previous results, intramuscular injection of AS1842856 reversed the LL2-induced changes of Deaf1, LC3, and ATG16L1, and PIK3C3, but did not affect LL2 tumor growth (Figure 10B-J and S7E-H), suggesting that FOXO-Deaf1-autophagy axis is critical for cancer-induced muscle wasting. Collectively, these findings indicate that dysregulated Deaf1 contributes to muscle regeneration defects during aging and cancer cachexia and may play a role in other myopathies.

**Figure 9.**
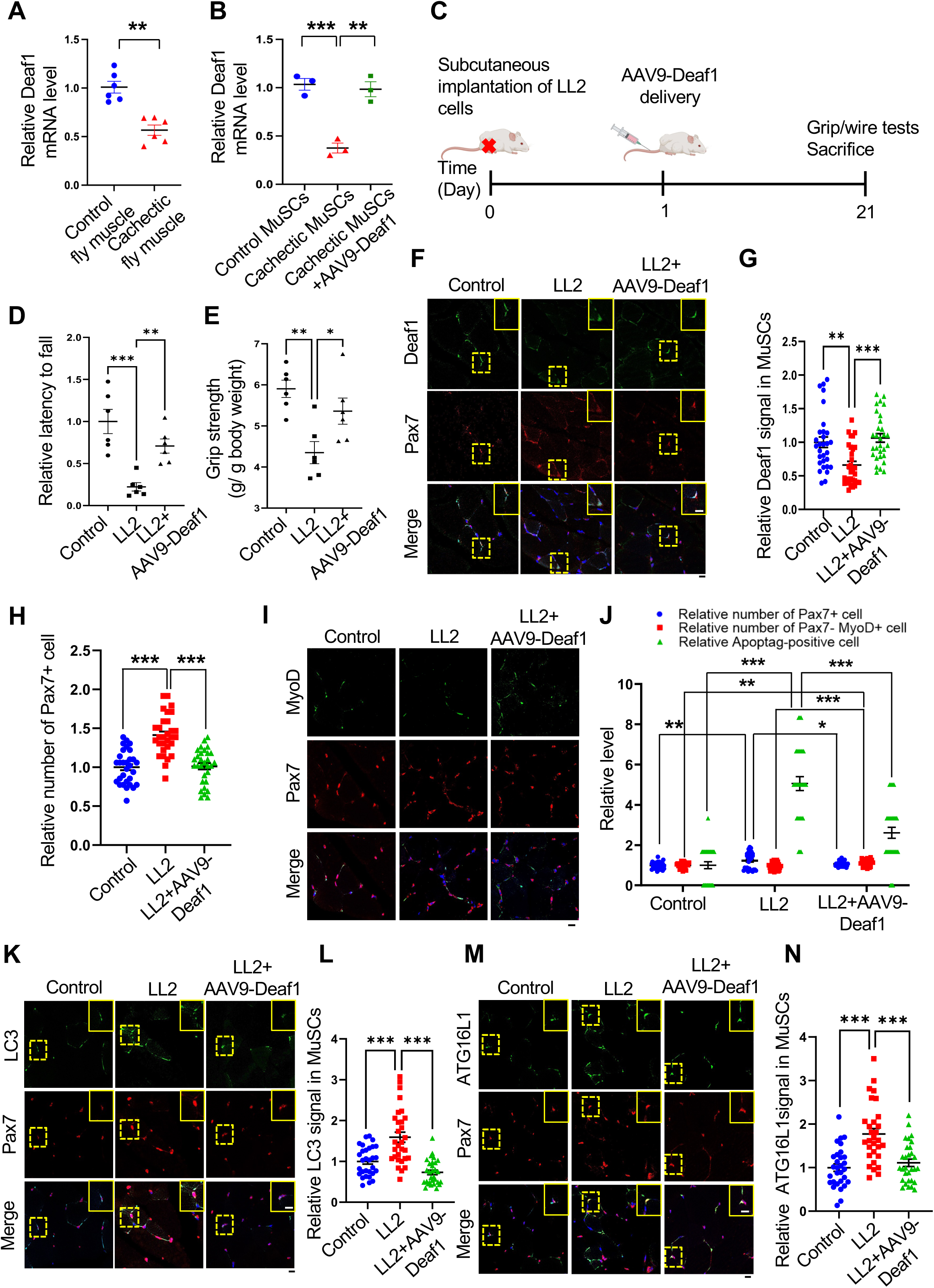
Enhanced *Deaf1* expression relieves muscle atrophy under cachectic conditions. (A-B) *Deaf1* mRNA is decreased in cachectic MuSCs. RNAs extracts from thorax of control (*esg-Gal4, UAS-GFP, tubGal80^ts^ or EGT*) or cachectic flies (*EGT; UAS-yki^3SA^*) (A) as well as from isolated MuSCs of NOD/SCID mice (∼3 months) with or without LL2 cachectic cell implantation, followed by AAV9-Deaf1 injection (B) were subjected to qPCR analysis. (C) Flow chart showing the details of mouse experiment using AAV9. (D-E) Augmented Deaf1 level increases muscle strength during cancer cachexia. Purified AAV9-Deaf1 viruses were systemically delivered using intravenous (IV) injection into NOD/SCID mice bearing LL2 tumors. After 21 days, muscle strength and function of young and aging mice was assessed by wire-hang (D) and grip strength (E) analyses. (F-N) An increase in Deaf1 expression represses autophagy and relieves muscle atrophy induced by cancer cachexia. Immunostaining of soleus muscles from NOD/SCID mice bearing LL2 tumors or LL2 tumors with AAV-Deaf1 IV injection using anti-Deaf1 (green) (F), anti-MyoD (green) (I), anti-LC3 (green) (K), anti-ATG16L1 (green) (M), and anti-Pax7 (red) antibodies. Deaf1 (G), Pax7 (H and J), Pax7-/MyoD+ (J), LC3 (L), and ATG16L1 (N) signals in MuSCs were quantified. Apoptotic cells were detected by ApopTag® In Situ Apoptosis Detection Kit and quantified in (J). Scale bar: 10 µm.

**Figure 10.**
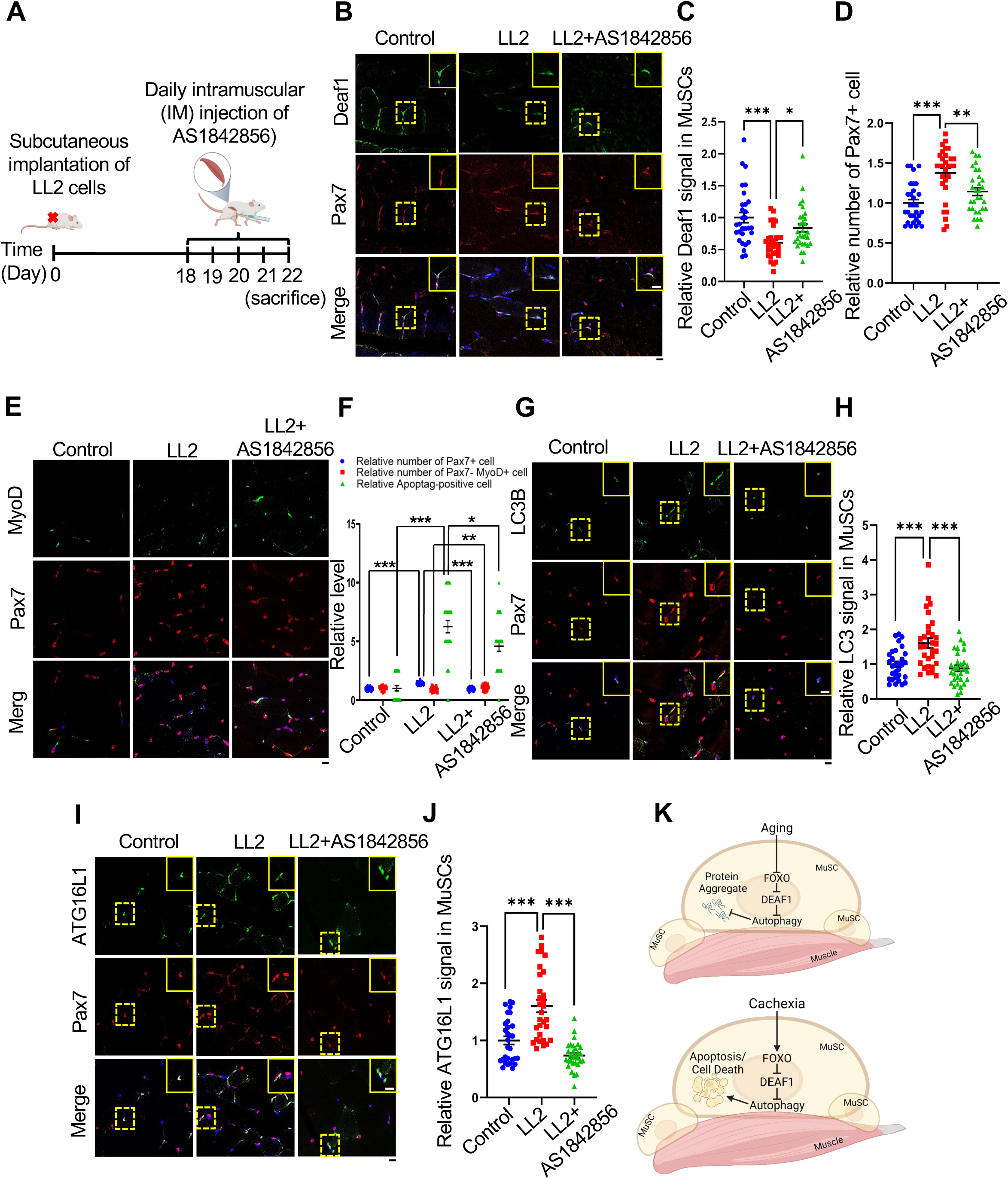
FOXO1 inactivation increases Deaf1 expression and improves muscle defects induced by cancer cachexia. (A) Schematic representation of the experimental design. (B-J) Immunostaining of soleus muscles from control mice or mice bearing LL2 tumors intramuscularly injected with a FOXO1 inhibitor, AS1842856, using anti-Deaf1 (green)(B), anti-MyoD (green)(E), anti-LC3 (green)(G), anti-ATG16L1 (green)(I), and anti-Pax7 (red) antibodies. Deaf1 (C), Pax7 (D and F), Pax7-/MyoD+ (F), LC3 (H), and ATG16L1 (J) signals in MuSCs were quantified. Apoptotic cells were detected by ApopTag® In Situ Apoptosis Detection Kit and quantified in (F). Scale bar: 10 µm. (K) Working model. Model showing that aging enhances *Deaf1* expression through FOXO signaling to transcriptionally suppress autophagy and in turn increase protein aggregates in MuSCs. In contrast, cachectic cancers activate FOXO to reduce *Deaf1* expression and promote autophagy, leading to cell death. Modulation of *Deaf1* levels can improve muscle atrophy induced by aging and cancer cachexia. Created with BioRender.com.

## Discussion

In this study, we identified the transcription factor Deaf1 as a novel regulator of muscle regeneration. We showed that Deaf1 functions downstream of FOXOs to regulate autophagy as well as to control MuSC renewal and differentiation. FOXOs repress *Deaf1* expression through binding to its promoter and reduced Deaf1 levels, which in turn increases the expression of multiple Atg genes, including *Atg16l1* and *PI3KC3*, thus activating autophagy. In contrast, Deaf1 suppresses expression of *Atg16l1* and *PI3KC3* and inhibits autophagy, promoting protein aggregate formation and cell death in MuSCs. Intriguingly, both upregulation and downregulation of Deaf1 led to defects in muscle regeneration. Moreover, changes in *Deaf1* expression levels were observed in aged and cachectic MuSCs. We further showed that pharmaceutical or Adeno-Associated Virus 9 (AAV9) targeting of the FOXOs-Deaf1-autophagy transcriptional cascade was able to improve muscle regeneration and functions under aging and cancer cachectic conditions (Figure 10K).

### Deaf1 is identified as a regulator of muscle regeneration from a *Drosophila* genetic screen

Multiple models were used to study MuSCs in attempts to treat muscle-degenerative diseases [52]. *Drosophila* has emerged as a new model to studying muscle regeneration due to the newly discovered MuSCs in adult flies [28-30]. The availability of numerous genetic tools makes *Drosophila* an indispensable and ideal model organism to investigate MuSCs. Our genetic screen in *Drosophila* utilized the cell lineage tracing system with a MuSC-specific *Zfh1* enhancer and identified a transcription factor, Deaf1, as a key regulator of muscle regeneration.

Multiple transcription factors have been implicated in a hierarchical transcription factor cascade required for muscle regeneration. For instance, MyoD, Myf5, myogenin, and MRF4, are myogenic regulatory factors that are upregulated during muscle regeneration and control MuSCs sequentially to differentiate into myogenic lineage cells [53]. In this study, we identified another transcriptional factor cascade, the FOXOs-Deaf1-autophagy axis. Upregulation or downregulation of this signaling axis results in MuSC cell death and suppression of muscle differentiation, suggesting that fine tuning of this signaling cascade is required for muscle regeneration.

### Deaf1-regulated autophagy in muscle regeneration

Increasing evidence demonstrates the importance of autophagy in the maintenance of MuSCs [54]. Autophagy-induced organelle and protein turnover is actively maintained in quiescent MuSCs [13]. Increased stress in senescent MuSCs, caused by suppression of autophagy, enhances susceptibility to apoptosis [55]. In this study, we also observed the same phenomenon. Deaf1 inhibits the expression of two autophagy regulators, ATG16l1 and PIK3C3. Thus, Deaf1 overexpression in *Drosophila* and mouse MuSCs leads to autophagy inhibition as well as decreases in MuSCs numbers and their differentiation. Recent studies on Atg16l1 and PIK3C3 further highlighted the need for autophagy in the maintenance of muscle mass and myofiber integrity under physiological conditions. For instance, Atg16l1 hypomorphic mice exhibited decreased levels of autophagy which resulted in a significant reduction in the generation and growth of muscle fibers [56]. Likewise, decreased levels of PIK3C3 were observed in the tibialis anterior muscles of older mice indicating a blockage of autophagy in aged muscles [57]. Our findings thus highlight Deaf1 as a critical regulator of autophagy in MuSCs and its functions in muscle regeneration. In our results, modulation of autophagy only partially rescued Deaf1-induced defects in MuSC renewal and differentiation, indicating that there might be other factors mediating Deaf1-dependent effects. Indeed, our Deaf1 ChIP-seq dataset revealed a significant enrichment of genes involved in transcription, translation, and stem cell population maintenance. Future studies will be required to investigate whether these genes regulate muscle regeneration.

### FOXOs-Deaf1 axis in the regulation of autophagy

FOXO (Forkhead box O) transcription factors, which are well conserved from C. elegans to humans, bind two consensus sequences, the DAF-16 family member-binding element (DBE) 5′-GTAAA(T/C)AA-3′ and the insulin responsive element (IRE) 5′-(C/A)(A/C)AAA(C/T)AA-3′ [46]. *Drosophila* only has one FOXO gene, while mammals have four FOXO genes: FOXO1, FOXO3, FOXO4, and FOXO6 [58]. We identified an IRE site in both *Drosophila* and mouse Deaf1 promoter regions. We further showed that FOXOs can bind to the promoter region of *Deaf1* and inhibit *Deaf1* expression, demonstrating that Deaf1 is a direct target of FOXOs in *Drosophila* and mouse. Our results thus suggest that the FOXOs-Deaf1 axis is evolutionarily conserved.

FOXOs are critical regulators of autophagy. It has been shown that FOXO can bind to genomic regions containing key autophagy regulators, Atg3 and Atg17, in muscles [43]. Despite its direct involvement in transcribing Atg genes, FOXOs suppress Deaf1 expression to activate autophagy by transcriptional upregulation of Atg16 and PI3KC3. Deaf1 overexpression can reverse FOXOs-induced autophagy, suggesting that Deaf1 mediates FOXOs-induced autophagy. Why do FOXOs need to regulate Deaf1 to activate autophagy? It is possible that FOXOs coordinates with Deaf1 to promote expression of all autophagy-related genes. As FOXO may only promote transcription of certain Atg genes, it requires additional autophagy regulators to induce autophagy [43]. Another possibility is that the FOXOs-Deaf1 axis can coordinate to modulate autophagy levels. Autophagy activity can be adjusted by the expression levels of Atg genes [17, 38, 59]. Thus, FOXOs may regulate additional autophagy regulators to further enhance Atg expression in response to different physiological stresses, with synergistic effects that can promote autophagy activities to a greater extent. Our findings thus suggest that FOXOs may regulate autophagy at multiple layers.

### Distinct mechanisms behind the same phenotypes of muscle diseases

Sarcopenia is the age-induced loss of muscle mass with diminished ability of muscle regeneration. The regenerative ability of skeletal muscles is also disturbed upon cancer cachexia, a syndrome characterized by the cancer-induced loss of skeletal muscle mass [3]. Although declines in MuSC activities have been reported in both muscle diseases, the underlying mechanisms remain unclear. We observed that *Deaf1* expression level was changed under sarcopenia or cancer cachexia conditions. To our surprise, *Deaf1* mRNA level is increased in aging MuSCs, but decreased in cachectic MuSCs, suggesting the different mechanisms underlying sarcopenia and cancer cachexia. Similar to Deaf1, the activities of FOXOs vary in different muscle diseases. Inactivation of FOXOs has been reported in aged muscles and MuSCs *[10, 43].* In contrast to sarcopenia, FOXOs expressions were increased in skeletal muscles under cancer cachectic conditions [24, 25]. Thus, our findings along with studies from other labs demonstrate that, even though muscle regeneration defects commonly occur in many muscle diseases, the molecular mechanisms behind these diseases could be different or even opposite.

At present, Deaf1 inhibitors or activators are unavailable. We thus utilized FOXO inhibitor/activator as well as adeno-associated virus 9 (AAV9) to manipulate Deaf1 expression levels. The intramuscular injection of a FOXO activator, LOM612, was able to restore *Deaf1* expression and MuSC numbers in aging muscles. Conversely, injection of a FOXO1 inhibitor, AS1842856, reversed the effects induced by cancer cachexia. Adeno-associated viruses have been shown to increase transduction efficiency and prolong stable gene expression. Among AAVs, AAV-serotype 9 (AAV9) can effectively transduce skeletal muscles and MuSCs [60, 61]. Enhanced *Deaf1* expression mediated by AAV9 alleviated muscle wasting induced by cachectic lung cancer, while *shDeaf1* expression restored muscle regeneration in aged soleus muscle. These two methods efficiently modulated Deaf1 expression levels in MuSCs, which potentially would aid in the development of new therapeutics against myopathies. Although our immunofluorescence data suggest that *Deaf1* is enriched in MuSCs, we also observed that FOXOs-Deaf1 axis occurs in C2C12 myocytes and other tissue cells, raising the possibility that alteration of Deaf1 levels in muscles contributes to the improvement of muscle atrophy. Future studies will be needed to examine the roles of muscular Deaf1 in muscle diseases.

## Materials and Methods

### Drosophila Husbandry

Flies were maintained in an incubator on 12-hour light/12-hour dark cycles, at 25°C with 60% humidity. The flies were fed with standard cornmeal/soy flour/yeast fly food. Fly stocks were obtained from the Bloomington Drosophila Stock Center (BDSC), FlyORF, and the Vienna Drosophila Resource Center (VDRC), as detailed in Table S3.

### RNAi Screening and Cell Quantification

Virgin females of TubGal80^ts^; Zfh1-Gal4, UAS-G-Trace muscle stem cell lineage tracing fly line were crossed with males of each RNAi line (8 females to 4 males), and the crosses were maintained at 18°C until eclosion of the F1 progenies. F1 expressing TubGal80^ts^, Zfh1-Gal4, UAS-G-Trace, UAS-RNAi were selected and maintained for 7 days at 29°C, with regular fresh food changes every 2 days, before dissection. Thoraxes were dissected from the F1 adult flies, fixed in 4% Paraformaldehyde (Electron Microscopy Sciences, 15710), before being cut into halves along the sagittal plane. The thoraces were then imaged under fluorescence microscopy (Olympus IX83) to quantify for MuSCs and their differentiated progenies.

### Immunostaining, Proteostat and Apoptag Staining, and Confocal Imaging

To monitor autophagy influx in the *Drosophila*, Chloroquine (Cayman, 14194) was added during the preparation of standard yeast fly food (10mg/ml), and flies were maintained on Chloroquine-containing fly food for 3 days at 25°C until dissection. Thorax muscle tissue harvested from *Drosophila* were fixed and cut into halves as described in the earlier section. The tissues were then washed, stained with Phalloidin at 1:1000 dilution, and incubated at 4°C overnight. The muscle tissues were then washed and stained with DAPI (1 ug/ml) for 10 min before mounting.

Cells were fixed with 4% Paraformaldehyde or iced methanol before blocking with 5% goat serum. The cells were then incubated with primary antibodies at 4°C overnight. Secondary antibodies were added and incubated for 1 h at room temperature, before DAPI (1 µg/ml) was added for 10 min. PROTEOSTAT^®^ Protein Aggregation Assay (Enzo, ENZ-51023) was used for detection of protein aggregates according to the manufacturer’s protocol.

Muscle tissues were harvested from sacrificed mice and fixed in 10% formalin (VWR, 11699404). The fixed tissues were dehydrated, embedded in paraffin, and subjected to sectioning. Briefly, the tissue sections were deparaffinized, antigen retrieved and blocked using PBS containing 1% BSA and 10% goat serum for 1 hr at room temperature. Primary antibodies were then added and incubated overnight at 4°C, followed by washing and incubating with a secondary antibody for 1 hour at room temperature. Tissue sections were then washed and mounted. For Apoptag staining, the muscle sections were deparaffinized, and stained with Apoptag® *In Situ* Apoptosis Detection Kit (Sigma-Aldrich, S7100) according to the manufacturer’s protocol.

Samples were visualized by Zeiss LSM 710 confocal microscope (ZEISS, Oberkochen, Germany). Antibodies used are documented in Table S3. For quantification, we quantified 30 images from 6 male mice per group (5 images per mouse).

### Cell Culture and Drug Treatments

C2C12 myoblasts were grown in DMEM (Sigma Aldrich, D5796) supplemented with 1% Penicillin-Streptomycin (Gibco, 15140122), and 10% Fetal Bovine Serum (FBS) (Sigma, F7524). DMEM supplemented with 2% Horse serum (Gibco, 16050-122) and 1% Penicillin-Streptomycin was given to C2C12 myoblasts to induce differentiation. For Bafilomycin-A1 (MedChemExpress, HY-100558), AS-1842856 (MedChemExpress, HY-100596) and LOM612 (MedChemExpress, HY-101035) treatments, C2C12 myoblasts were treated with or without 0.1 µM Bafilomycin-A1, 50 µM AS-1842856 or 10 µM LOM612, with corresponding volume of DMSO as control for 2 days.

### Muscle Stem Cell Isolation

Muscle stem cells (MuSCs) were isolated from the hindlimbs of mice. Briefly, the muscle tissues were first dissected and collected in DMEM, before mincing into a slurry. Subsequently, they were digested in 1.5 mg/ml Collagenase/Dispase (Roche, 11097113001) dissolved in DMEM, containing 1% Penicillin-Streptomycin and incubated at 37°C with gentle agitation at 600 rpm, for 1 hour. The minced muscle tissue was triturated before passing through a 70 µm nylon mesh strainer and 40 µm nylon mesh strainer. The flow-through was centrifuged at 1000 g for 10 min. The cell pellet was then resuspended in MuSC growth medium (DMEM, 20% FBS, 10% Horse serum, 1% Chick Embryo Extract (Life Science Production, MD004A-UK), bFGF (5 ng/ml) (Gibco, PHG0021), 10 µM ROCK inhibitor (MedChemExpress, HY119937), 1% sodium pyruvate (Lonza, BE13-115E) and 1% Penicillin-Streptomycin). The cells were pre-plated overnight. The supernatant containing unattached cells were then transferred to a pre-coated Gelatin (Sigma-Aldrich, G6650) plate to be grown and cultured.

### Plasmids

The full-length of mouse Deaf1 cDNA was amplified and cloned from our C2C12 cDNA library, with primers used as detailed in Table S3, into pLX313 backbone plasmid (Addgene, #118014). Constitutively active FoxO1-ADA-GFP (Addgene, #35640) and FOXO3-AAA (Addgene, #1788) were obtained from Addgene. To knockout *Deaf1*, we utilised CRISPR as previously described [62, 63] to silence the expression of *Deaf1* in C2C12 myoblasts, with sgRNA sequences documented in Table S3. To knockdown the expressions of Deaf1, FOXO1, and FOXO3, shRNAs with sequences listed in Table S3 were cloned into tet-inducible vector Tet-pLKO-puro (Addgene #21915). To generate luciferase reporters, the Deaf1 promoter region was first PCR-amplified from genomic DNAs of C2C12 mouse myoblasts and cloned into the pGL4.20 vector (Promega, E675A)., with primers detailed in Table S3, using KOD One PCR Master Mix (Toyobo, KMM-101). Using PCR site-directed mutagenesis, we deleted the FOXO consensus sequence within the mouse Deaf1 Promoter using primers as shown in Table S3.

### Construction and generation of AAV9-Deaf1 and AAV9-shDeaf1

Deaf1 cDNA was cloned into pAAV-CAG-GFP (Addgene, #37825), and Deaf1 shRNAs were cloned into pAAV-Ptet-RFP-shR-rtTA (Addgene, #35625). The Deaf1 and Deaf1-shRNA carrier AAV vectors were transfected with pAAV-R/C (AAV serotype 9 [AAV9]) and an adenoviral helper vector in HEK293T cells. Viruses were purified with AAVpro Purification Kit Maxi (All Serotypes) (Takara, cat# 6666), as per manufacturer’s protocol. Following that, the titers of the generated AAV vectors were determined by Takara-AAVpro Titration Kit (for Real Time PCR), Ver.2 (Takara, cat# 6233), as per manufacturer’s protocol.

### Chromatin immunoprecipitation (ChIP)

Chromatin Immunoprecipitation was performed on thorax muscle tissues of FOXO-V5;FOXO^25^/ FOXO^25^ *Drosophila* (BDSC, 80945) or C2C12 cells, using the Simple ChIP Plus Enzymatic Chromatin IP Kit (Magnetic Beads) (Cell Signaling Technologies, #9005), according to manufacturer’s instructions and following the procedures described in our previous paper [18]. Briefly, the C2C12 cells and grinded muscles from *FOXO-V5; FOXO^25^/FOXO^25^* flies were crosslinked with 1% Formaldehyde and quenched with glycine. DNA-co-immunoprecipitations with IgG control antibody (a component of the kit), anti-Deaf1 (Bethyl Laboratories, #A303-187A), anti-FOXO1 (Cell Signaling Technologies, 2880S), anti-FOXO3 (Santa Cruze, sc-48348), or anti-V5 antibody (Biorad, MCA2894), were analyzed by deep DNA-sequencing or quantified by ChIP-qPCR using primers listed in Table S3.

For ChIP-seq data analysis, quality control was performed using FastQC (v0.11.8) and ChIPQC (1.32.2). ChIP-seq reads were aligned to the UCSC mm10 genome using bwa (0.7.17-r1998-dirty) [64]. Duplicates and unmapped reads were flagged with samblaster (0.1.24) and removed with sambamba (0.7.0) [65]. Peaks were called using macs2 (2.1.2) with the significance cut-off q-value at <= 0.1 [66]. Peaks were merged and filtered using the idr R package. Only highly reproducible peaks with IDR between 0 and 0.05 (inclusive) were kept.

### RNA-seq analysis

RNA-seq data for cancer cachexia was downloaded from GEO – (GSE133979) [67] and processed using STAR [68] and RSEM [69]. RNA-seq data investigating sarcopenia in human patients was downloaded from GEO (GSE111010 and GSE111016) [70]. Differential expression analysis was performed using DESeq2 [71].

### Luciferase Assay

C2C12 myoblasts were transiently transfected using Lipofectamine 3000 Transfection Reagent (Invitrogen, 100022052), with Deaf1-promoter-pGL4.20, Deaf1-promoter^ΔIRE^-pGL4.20 and Renilla luciferase control. Two days after transfection, cells were treated with or without 10 µM LOM612 (MedChemExpress, HY-101035) for 5 hrs and harvested for luciferase assay. Dual Luciferase reporter assay (Promega, E1960) was performed according to manufacturer’s protocol, with luciferase activity measured by Tecan Infinite M200 Plate Reader, using Renilla luciferase activity as the internal control.

### Mouse and Ethical Approval

Mice were housed in individually ventilated cages under 12-hour light/12-hour dark cycles, with food and water ad libitum. Experiments were carried out in Duke-NUS Medical School, Singapore, according to ethics approval by the Institutional Animal Care and Use Committee (IACUC) of Duke-NUS Medical School, Singapore. For all *in vivo* work in mice, we used 6 male mice per group for all experiments.

Young C57BL/6J mice used were 3 months old and old C57BL/6J mice used were at least 18 months old. Mice with intravenous injection of AAV9-shDeaf1 at 1×10^10^ viral particles or with intramuscular injection of LOM612 at concentration of 2 µM into hindlimb of old mice every other day for a total of 10 injections, were subjected to grip strength test and wire hanging test at the end of experiment before sacrificing. AAV9-shLuc or DMSO (controls) was used to inject young and old mice.

For the mouse cachexia model, 3×10^5^ of LL2 cells were implanted subcutaneously on NSG mice aged 6-8-week-old. After LL2 implantation, AAV9-Deaf1 was injected intravenously at 1×10^10^ viral particles/ site of injection or with PBS. Alternatively, 18 days after LL2 implantation, the mice were injected intramuscularly with 10 µl of 50 µM AS1842856, into the hindlimbs of the mice. The injections were made once a day, consecutively for 5 days, before sacrifice. Tumor growths were measured throughout the experiment. Grip strength test and wire hanging test were performed at the end of experiment before sacrificing.

### Mouse Cryoinjury Treatment

The mice were first anesthetized with isoflurane and the hindlimb was applied with betadine for sterilization. An incision was made on the skin to expose muscles. A metal probe with diameter of 2 mm was immersed in liquid nitrogen before being applied to the muscles for 10 seconds, twice. The skin was then sutured and held in place with Histoacryl^®^ (Braun). Betadine was applied to the wound and the animal was allowed to recover. The animals were sacrificed after 2 weeks, and muscles were harvested.

### Mouse Grip Strength Test

Muscle strength was tested with a grip strength test. The grip strength of the mouse was measured with a grip strength meter (Ametek, DFE-II) mounted horizontally with a non-flexible grid. Each mouse was allowed to grasp the grid with all four paws or only the front two paws and was pulled horizontally until its grasp was broken. The force in grams per pull was recorded. The hindlimb grip strength was calculated by subtracting the grip strength of only the front paws from the grip strength of all four paws, and the grip strength readout is defined as the gram force divided by mouse body weight.

### Mouse Inverted Screen Hanging Test

The inverted screen hanging test was used to assess muscle strength and endurance. Before the test, the mice were weighed, and their masses were recorded. The mice were then placed on a wire grid surrounded by a wooden frame before being inverted gently and held 50 cm above a soft, padded surface. The time that the mouse held onto the wire grid was measured until the mouse released its grip and fell. The inverted screen hanging test was repeated thrice, with a resting period between each replicate, and the average of the triplicates was calculated. The “holding impulse” was used as an output measure, which was the average time (s) that the mouse held on before falling over divided by mouse body weight.

### RNA Isolation and qRT-PCR

Total RNAs were extracted from cultured cells or tissues using TRIzol reagent (Invitrogen, 15596-018) according to manufacturer’s protocol and were reverse transcribed to cDNA with PureNA First Strand cDNA Synthesis Kit (Research Instruments, KR01-100). qRT-PCR was performed using BlitzAmp Hotstart qPCR Master Mix (Mirexes, 1204201) with CFX96 Real-Time System (BIO-RAD, Hercules, CA, USA) as per manufacturers’ protocols. The expression levels of genes on interests were analyzed using Bio-RAD CFX Manager Software, with Actin and rp49 as internal controls. Each qRT-PCR was performed in technical triplicates, and primer sequences used are listed in Table S3.

### Statistical Analysis

Results are expressed as mean ± SEM. Student’s t-test (two sample comparisons) or one-way ANOVA (multiple sample comparisons) analysis was performed with GraphPad Prism to determine the significance. Differences were considered significant if p values were less than 0.05 (*), 0.01 (**), 0.001 (***). Analysis of Deaf1 ChIP-seq and transcriptomic datasets was performed using R/Biconductor.

## Data and code availability

ChIP-seq data has been submitted to GEO (GSE237088). Code to recreate the reported ChIP-seq and transcriptomic analyses are available from https://github.com/harmstonlab/Goh_et_al_deaf1.

## Acknowledgements

We thank the Vienna Drosophila Resource Center, FlyORF, and the Bloomington Drosophila Stock Center for fly stocks. We also thank members of the Tang laboratory for experimental help and critical comments on the manuscript. This work was supported by Singapore’s Ministry of Education AcRF Award (2022-MOET1-0004, to H.-W.T.; MOE2019-T2-2-006, to N.Y.F; and MOE-T2EP30121-0013, to N.Y.F), National Academy of Medicine grant (MOH-001189-00, to H.-W.T.), National Medical Research Council (NMRC) (MOH-001208-00 to H.-W.T.; MOH-OFIRG20nov-0018 to N.Y.F; MOH-OFYIRG19nov-0022 to H.S.C), National Taiwan University Hospital (112-L2003 to S.-Y.H), the Singapore National Research Foundation (NRF-NRFF12-2020-0008 to J.N), and National University of Singapore and Yale-NUS College (through Reimagine Research Grant IG20-RRSG-001 to N.H.).

